# Balancing disease control efficacy, resistance durability, and economic outcomes: Insights from a participatory modeling study integrating grapevine cultivar deployment timing with fungicide treatments

**DOI:** 10.1101/2025.09.04.674243

**Authors:** Marta Zaffaroni, Adeline Alonso Ugaglia, Anne-Sophie Miclot, Loup Rimbaud, Julien Papaïx, Jean-François Rey, Frédéric Fabre

**Affiliations:** INRAE, Bordeaux Sciences Agro, SAVE, 33882 Villenave d’Ornon, France; INRAE, BioSP, 84914 Avignon, France; INRAE, Pathologie Végétale, 84140 Montfavet, France

**Keywords:** resistant cultivars, resistance durability, downy mildew, companion modeling, grapevine mathematical modeling, vine-growing territory, cost-benefit analysis, coconstruction workshops, wine cooperative

## Abstract

In this study, we developed, assessed and compared strategies for deploying disease-resistant grapevine cultivars in partnership with a wine cooperative in South-West France. In four collaborative workshops, cooperative managers and multidisciplinary researchers jointly designed six feasible deployment strategies. These scenarios, implemented within real vineyards, involve planting resistant cultivars (RC) in vineyards (i) more than 30 years old, (ii) on 3.3% of the older plots each year or (iii) on plots located in no-treatment zone (due to proximity to watercourses or human dwellings), with and without a fixed maximum percentage of resistant cultivars in the vineyard. The strategies were evaluated with the *landsepi* model, which simulates the spread and evolution of pathogens across agricultural landscapes in response to host resistance deployment. The assessment criteria included downy mildew control efficacy, resistance durability, decrease in fungicide use, and economic benefits for individual vineyards and the cooperative. These scenarios led us to study more specifically the effects of two little-studied factors: (i) the presence or absence of fungicide applications and (ii) the massive or progressive introduction of resistant cultivars. Fungicide application substantially decreases the risk of resistance breakdown and complements the epidemiological control provided by resistant cultivars, particularly when the fitness costs of virulence are low. The probability of pathogens adapted to RC becoming established was generally lower with progressive than massive RC introduction; however, when establishment did occur, the time to establishment was shorter for progressive strategies at a cropping ratio ≥ 50%. Ultimately, the choice of RC introduction strategy had a minimal overall impact on disease control due to compensatory epidemiological and evolutionary processes, including dilution effects, and the probability and timing of resistance breakdown. These results are discussed in the broader framework of action-research approaches and the challenges and opportunities for resistant cultivar adoption by vine-growers. Our study highlights the potential of participatory approaches for developing practical, scientifically grounded strategies for sustainable viticulture.

## 2 Introduction

Growing cultivars resistant to plant disease is a relatively low-input and cost-efficient strategy for protecting agricultural crops against pathogens [1]. This approach, used to manage diseases in diverse crops, depends on lengthy and costly breeding processes, especially for perennial crops. In grapevine breeding, the selection of resistant rootstocks was effective against phylloxera, but the hybrids developed in the late 19th century have been less successful. These hybrids were initially created with resistance genes originating from American wild *Vitis* species in response to the devastation of European vineyards by mildews [2]. The hopes for the replanting of European vineyards initially raised by these hybrids were dashed during the first half of the 20th century, principally due to their poor enological qualities. With the discovery of the effect of copper on grapevine downy mildew (GDM) in 1882 [3] and the development of synthetic pesticides after World War II, the protection of grapevines against disease was based almost exclusively on the use of fungicides. The mean treatment frequency index (TFI, [4]) for French vineyards over a 10-year period (2010-2019) is currently 12.1, with fungicides accounting for 83% to 86% of the total TFI [5]. Fungicides are applied mostly to control GDM and powdery mildew, the two most important diseases of grapevine in France [6].

The current paradigm of grapevine protection dominated by fungicide use may change in the future with the planting of mildew-resistant cultivars. Since the early 1980s, European breeding programs focusing on mildew resistance have produced several resistant cultivars [7]. Growers can currently manage GDM, the target disease of this study, by planting resistant cultivars (RC) bearing a single resistance gene, principally *Rpv1, Rpv3*.*1, Rpv10* or *Rpv12*, or pyramided RC, combining both *Rpv1* and *Rpv3*.*1*. These pyramided cultivars were developed in the INRA-ResDur breeding program [8]. The breeding of pyramided cultivars is more costly and time-consuming, but is often preferred by breeders as a means of increasing the durability of resistance [9]. Indeed, a significant challenge in the breeding of RC is that pathogens frequently evolve rapidly, leading to resistance breakdown very soon after deployment, sometimes resulting in catastrophic epidemics [1, 10, 11].

Over the last two decades, the focus has shifted from the search for durable resistance [12] to the durable management of resistance. Durability is no longer seen as an intrinsic property of resistance genes but as the outcome of strategies incorporating the development of resistant cultivars by breeders and their spatial and temporal deployment by farmers [13]. Deployment strategies should be compared in terms of their efficacy (*i*.*e*. provision of satisfactory epidemiological control) and durability (*i*.*e*. provision of satisfactory evolutionary control despite the evolutionary potential of the pathogen) [14]. Experimental testing of these strategies is challenging, and often impossible, due to large number of combinations of options for the deployment of resistance and the large spatiotemporal scales involved. Mathematical models have been developed to compare and potentially optimize deployment strategies (for a review, see [13]). A recent study [15] using the spatially explicit stochastic model *landsepi*, showed that the evolutionary control provided by a cultivar pyramiding *Rpv1* and *Rpv3*.*1* is placed at risk when a single-gene RC carrying one of these genes is planted concomitantly in the landscape, particularly if the pathogen has a low probability of mutation. These results were obtained by exploring a wide range of possible cropping ratios (*i*.*e*. the proportion of candidate fields sown with the RC) in theoretical agricultural landscapes consisting of uniform square fields.

As in almost all studies based on mathematical models, the model *landsepi* used in this last study [15] was used to compare top-down deployment strategies, regardless of the perception of farmers. By contrast, here, we compare bottom-up strategies identified during four action-research workshops bringing together professional stakeholders and researchers from different scientific disciplines. Action-research is an effective approach for the design of sustainable farming systems through the explicit integration of both scientific and local knowledge and practices [16]. It frequently uses process-based modeling as a tool for understanding, discussing, and exploring complex issues [17]. This action-research took place in spring 2023 and involved a wine cooperative located in south-western France. Right from the first workshop, it became clear that improvements were required in the representation of fungicide treatments and key aspects of wine economics in the *landsepi* model before the scenarios could be evaluated correctly. The deployment strategies identified with local stakeholders were then compared and their epidemiological, evolutionary, environmental and economic outputs were discussed. Finally, the bottom-up scenarios developed led us to study more specifically the effects of two factors little explored in previous studies despite their practical interest: (i) fungicide applications according to a severity threshold and (ii) the massive *versus* progressive introduction of RC.

## 3 Materials and methods

### 3.1 The Buzet cooperative and its territory

The Buzet Protected Designation of Origin (PDO) area, established in 1973, is located around the district of Buzet-sur-Baïse (coordinates: 44° 15 32 N, 0° 18 00 E; South-West France). Bounded by the River Garonne to the north and east, and by the Landes forest to the west, its territory extends over 27 municipalities. The PDO is an official European label ensuring the quality and origin of products. For the PDO label to be awarded, specific production rules (planting density, list of grape cultivars that can be planted, maximum yield and maximum number of missing plants) must be respected. During the research project, a partnership was established with the “Nous, les vignerons de Buzet” wine cooperative located in this area. A wine cooperative is an agricultural organization that produces and sells wine made from the grapes of its members. It performs joint wine-making, storage, packaging and sales operations on behalf of its members. Farmers very often delegate technical decisions to the cooperative advisors (diverse services including crop protection and choice of the cultivars to plant) [18]. This wine cooperative, founded in 1953, includes most of the producers of PDO wine in the area (160 producers in 2023), 1935 hectares under vines, and accounts for 95% of the output for this PDO. In 2020, production was about 13 million bottles. This cooperative lays down specifications to ensure a degree of uniformity in agricultural practices and production costs and a common policy for product promotions. It also provides grape growers with technical support for various aspects including the renewal of vines in their vineyards.

### 3.2 Identification of the resistant cultivar deployment strategies to be simulated

The planting of certain resistant grape cultivars on an experimental basis within PDO areas has been authorized since 2021 under the VIFA (in French, “Variétés d’Intérêt à des Fins d’Adaptation”, literally “Cultivars of Interest for Adaptation Purposes”) directive. The Buzet cooperative has requested such authorization for the INRAE Resdur cultivars Artaban and Vidoc (RC for the production of red wine). This authorization is valid for 10 years, with planting limited to a maximum of 5% of the surface area of the vineyard and 10% of the wine blend. These cultivars have not yet been planted in this PDO area.

The main objective of this work was to construct and compare strategies for the deployment of resistant cultivars on the cooperative’s territory. Collective scenarios for the deployment of GDM-resistant vine cultivars were coconstructed with team members from the wine cooperative, through four workshops organized on a monthly basis (January-June 2023) at the premises of the cooperative. The workshops were attended by the director of the wine cooperative, managers from the various departments of the cooperative (vineyard, marketing, research and innovation), and researchers from the INRAE team (epidemiology, biomathematical modeling, plant pathology, economics), under the eye of an independent sociologist acting as an observer, and in the presence of an illustrator to facilitate graphic representation. The workshops were organized by the ecodesign consultancy Think+, one of the cooperative’s partners for the definition of its sustainable development strategy.

The theme of the first workshop was “Understanding the issues and sharing objectives”. The *landsepi* model was presented as an educational tool for raising the awareness of cooperative staff about the challenges of RC deployment. During the second workshop, facilitation approaches enabled the team from the cooperative winery to identify a set of 12 exploratory deployment scenarios for simulation on the cooperative’s territory. These scenarios corresponded to the strategic objectives of the cooperative’s stakeholders. At the third workshop, the results obtained for the exploratory scenarios were presented, compared and discussed in terms of their ability to control GDM, promote the durability of resistance and reduce the number of fungicide treatments, and the economic benefits to both individual vineyards and the wine cooperative were calculated. This stage made it possible to eliminate several scenarios and to refine others. Six preferred scenarios were selected for testing and were compared with the status quo, in which plots continue to be replanted exclusively with the susceptible cultivars (SC) traditionally grown in the PDO area. The fourth and final workshop provided an opportunity to discuss the six selected scenarios in greater depth, to focus on the obstacles and levers to the implementation of one of these scenarios in the near future from the standpoints of different stakeholders (vine growers, consumers and wine cooperative), and to summarize the findings.

The *landsepi* model (see below) was used to compare the deployment strategies identified during the first and third workshops. It was computationally difficult to run the model simulation over the entire acreage of the Buzet cooperative. Working with the staff of the Buzet cooperative, we therefore selected two subdomains representative of the entire territory (Fig. 1A). The “Central” vineyard, at the core of the territory of the cooperative, is characterized by a high density of vines (21% of the surface area bounded by the perimeter of this subdomain is under grapevines). The “Diversified” vineyard is characterized by vineyard plots that are more scattered and located among other crops (2% of the surface area of this subdomain under grapevines). The main characteristics of the two subdomains are reported in Table 1.

**Table 1.** Details on the representative “Central” and “Diversified” vineyard subdomains. The mean and standard deviation of the attributes are reported when relevant.

| Description | Central vineyard | Diversified vineyard |
| --- | --- | --- |
| Number of fields | 1422 | 709 |
| Number of farmers | 57 | 36 |
| Field surface area (ha) | 0.43 (± 0.41) | 0.56 (± 0.58) |
| Field age (years) | 33.40 (± 15.10) | 34.77 (± 13.51) |
| Pairwise distance between fields (m) | 2598 (± 1329) | 5296 (± 2876) |
| Probability of field autoinfection<sup>a</sup> | 0.68 (± 0.12) | 0.70 (± 0.12) |
| Probability of field autoinfection and alloinfection<sup>b</sup> | 0.86 (± 0.07) | 0.86 (± 0.07) |

<sup>a</sup> Probability of an infectious propagule emitted in a given field dispersing within this field. <sup>b</sup> Probability of an infectious propagule emitted in a given field dispersing within this field or to any other field in the landscape.

**Fig 1.**
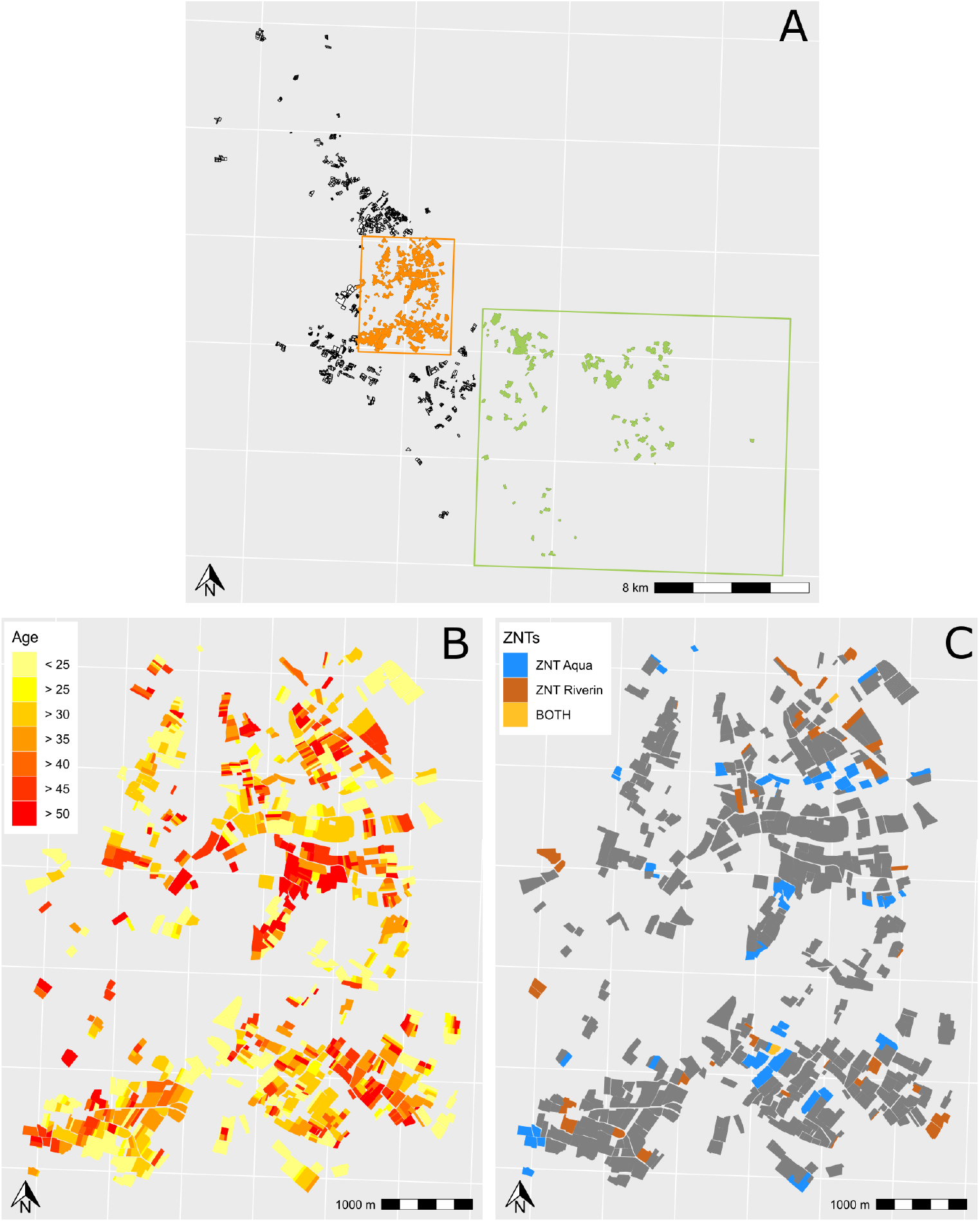
Maps of the Buzet vineyard. A: Two landscapes were considered in the Buzet territory. The “Central” vineyard corresponds to the central part of the Buzet area (in orange). This landscape (4.8 km × 6.0 km) includes 1422 fields and 57 vine-growers, with a mean field size of 0.43 ha. The “Diversified” vineyard corresponds to a peripheral part of the Buzet area (in green). This landscape (13.3 km × 16.0 km) includes 709 fields and 36 vine-growers, with a mean field size of 0.56 ha. B: Age of the plantations in the “Central” vineyard. C: Location of the fields in the “central vineyard” located in the no-treatment zone (NTZ) due to their proximity to the aquatic zone (“NTZ Aqua”) or a residential area (“NTZ Riverain”) or both. The maps for the “Diversified” vineyard are shown in Fig. S1.

### 3.3 Model description

#### 3.3.1 Model overview

We used the version of the model described by [15, 19]. It simulates the spread and evolution of a pathogen alternating within-season clonal reproduction and between-season sexual reproduction in a heterogeneous agricultural landscape over multiple cropping seasons. The demogenetic dynamics of the host-pathogen interaction is described by a HLIR structure (“healthy-latent-infectious-removed”). Hereafter, *H*_*i,v,t*_, *L*_*i,v,p,t*_, *I*_*i,v,p,t*_, *R*_*i,v,p,t*_, and *Pr*_*i,p,t*_ denote the numbers of healthy, latently infected, infectious and removed individuals and pathogen propagules in field *I* (*i*=1,…,*J*), for cultivar *v* (*v*=1,…,*V*), pathogen genotype *p* (*p*=1,…,*P*) at time step *t* (*t*=1,…,*T* × *Y*), where *T* is the total number of time steps in a cropping season *y* (*y* = 1, …, *Y*). In this model, an “individual” is defined as a given portion of plant tissue that can be infected by a single propagule, therefore excluding co-infection of the same part of the plant tissue by multiple propagules. An “individual” is referred to hereafter as a “host” for the sake of simplicity. As the host is cultivated, we assume that there is no host reproduction, dispersal or natural mortality (leaf senescence near the end of the cropping season is considered to be part of the host harvest). At the start of the first cropping season, susceptible hosts are contaminated with a primary inoculum consisting exclusively of non-adapted pathogens (denoted “WT” here for “wild-type”). At the start of subsequent cropping seasons, healthy hosts are contaminated with the primary inoculum generated by a single annual sexual reproduction event at the end of the previous cropping season. The propagules generated by sexual reproduction are gradually released during the following cropping season. Therefore, within a cropping season, healthy hosts may be infected by both the primary inoculum and by the secondary inoculum produced by infectious hosts through multiple clonal reproduction events. During the simulation, a WT pathogen can acquire an infectivity gene through a single mutation, or through sexual reproduction with another pathogen carrying such a gene. With two resistance genes, the single mutants “SM1” and “SM2” can break down the resistance conferred by the first and second resistance genes, respectively, and the superpathogen “SP” (with both infectivity genes) can break down the resistance conferred by both resistance genes (thereby overcoming the resistance of the pyramided cultivar). However, infectivity may be acquired with a cost to fitness*θ* [20, 21], *which, in our model, is associated with a lower probability of infection. Based on the values reported in previous studies* [15, 19] *for fitness costs θ* ≤ 0.25, we assumed that a fitness cost is paid only for unnecessary virulence (as opposed to a fitness cost paid on all hosts). The resulting host-pathogen interaction matrix for four cultivars (one SC, two RC carrying a single resistance gene and one RC carrying both genes) and the corresponding four pathogen genotypes is reported in Table 2.

**Table 2.** Host-pathogen interaction matrix

| | | Host genotype $v$ | | | |
| --- | --- | --- | --- | --- | --- |
|  |  | SC | RC<sub>1</sub> | RC<sub>2</sub> | RC<sub>12</sub> |
| Pathogen genotypes $p$ | WT | 1 | 0 | 0 | 0 |
| | SM<sub>1</sub> | $1-\theta$ | 1 | 0 | 0 |
| | SM<sub>2</sub> | $1-\theta$ | 0 | 1 | 0 |
| | SP | $(1-\theta)^2$ | $1-\theta$ | $1-\theta$ | 1 |

The matrix provides the coefficient by which the infection probability is multiplied. The value of this coefficient reflects the relative probabilities of infection for the wild-type (WT) and adapted (single mutants SM<sub>1</sub> and SM<sub>2</sub>, and superpathogen (SP)) pathogen genotypes on susceptible (SC) and resistant (RC) cultivars carrying a single major resistance gene (RC<sub>1</sub> and RC<sub>2</sub>), or both resistance genes (RC<sub>12</sub>). The fitness cost of infectivity $\theta$ , defined with respect to the major resistance genes considered, is paid only for unnecessary virulence (as opposed to fitness costs paid on all hosts).

Model equations are presented in *Supporting Information*, and the code is available from the R package *landsepi* (v1.3.0, [22]). With respect to the model previously described [15, 19], we added the possibility of applying fungicides to control the pathogen population. The corresponding modification of the model is described in section 3.3.2.

#### 3.3.2 Fungicide application

The effect of fungicides, the main method of GDM control, was incorporated as follows. We modified a model that can be parameterized for many fungicides and diseases to fit the case of a non-penetrant (*i*.*e*. contact) fungicide with pre-infection activity [29]. Specifically, we assumed that fungicide application reduces pathogen infection rate *e*_*max*_ as *e*(*t*) = *e*_*max*_ × (1 − *EF* (*t*)) where the fungicide effect *EF* (*t*) was given by:

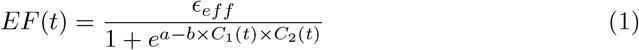

where *ϵ*_*eff*_ is the maximum fungicide efficiency, which coincides with fungicide efficacy at the date of application, *a* and *b* shape parameters and *C*_1_(*t*) × *C*_2_(*t*) is the concentration of fungicide at time *t*.

The fungicide is applied at the dose recommended. Its initial concentration is therefore equal to one, to ensure uniform coverage of the host tissue. As the fungicide is non-penetrant, its concentration decreases after application through both natural degradation and dilution by plant growth, as the new plant biomass is not protected by the contact fungicide. Therefore, 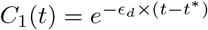 accounts for fungicide degradation, at a rate *ϵ*_*d*_, from the date of the last fungicide application *t*^∗^. The term 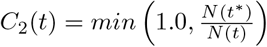 accounts for the newly produced host biomass, which is not protected by the fungicide. Note that *N* (*x*) = *H*(*x*) + *L*(*x*) + *I*(*x*) + *R*(*x*) is the total biomass of a given cultivar, in a given field, at time *x* = *t* or at the date of fungicide application *x* = *t*^∗^.

Fungicides can be applied on both SC and RC, on a set of potential treatment dates *t*^∗^ = 1, 4, 7, …., 109 during each cropping season. Treatments can be applied every three days, from the start of the simulation *t*^∗^ = 1 and ending at *t*^∗^ = 109 to comply with the respect of a preharvest period to prevent the presence of residues on grapes. At each of these dates, fungicides are applied in a given field if the severity of infection in this field exceeds a “fungicide application threshold” *ϵ*_*t*_. We therefore assumed that fungicides were applied if 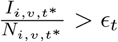

#### 3.3.3 Model parameterization for *Plasmopara viticola*

We used the parameterization proposed by [15, 19] to simulate epidemics of GDM caused by *Plasmopara viticola*, in fields sown with SCs or pyramided RC carrying the major resistance genes *Rpv1* and *Rpv3*.*1*. For fungicide treatment, we set the parameter values to ensure a total number of fungicide treatments by field and by season close to 10 in a landscape entirely populated by SC. This value is consistent with the number of fungicide treatments currently applied in French vineyards to control GDM and powdery mildew [5]. Unfortunately, no data were available to determine how many fungicide treatments were applied to control GDM only. All the model parameters used in the simulations are listed in Table 3.

**Table 3.** Summary of model parameters

| Notation | Parameter | Value | Source |
| --- | --- | --- | --- |
| Simulation Factors |  |  |  |
| $Y$ | Number of cropping seasons | 31 years | Fixed |
| $T$ | Number of time steps in a cropping season | 120 days | Fixed |
| $V$ | Number of host cultivars | 2 | Fixed |
| $J$ | Number of fields in the landscape | [1422; 709; 400]<sup>a</sup> | Varied |
| $P$ | Number of pathogen genotypes | 4 | Fixed |
| Initial conditions and seasonality (same value for all cultivars) |  |  |  |
| $C_v^0$ | Plantation density of host cultivar $v$ | 1 m<sup>-2</sup> | Fixed |
| $C_v^{max}$ | Maximal density of host cultivar $v$ | 20 m<sup>-2</sup> | Fixed |
| $\delta_v$ | Growth rate of host cultivar $v$ | 0.1 day<sup>-1</sup> | [23] |
| $\Phi$ | Initial probability of susceptible host infection | 5.10<sup>-4</sup> | Fixed |
| $\lambda$ | Probability of off-season survival for pathogen spores | 10<sup>-4</sup> | Fixed |
| Pathogen aggressiveness components |  |  |  |
| $e_{max}$ | Expected infection probability in a compatible host-pathogen interaction | 0.9 | [24, 25] |
| $\Gamma_{exp}$ | Expected duration of the latency period | 7 days | [19] |
| $\Gamma_{var}$ | Variance of latency period duration | 8 days | [19] |
| $\Upsilon_{exp}$ | Expected duration of the infectious period | 14 days | [19] |
| $\Upsilon_{var}$ | Variance of infectious period duration | 22 days | [19] |
| $r_{exp}$ | Expected propagule production rate | 2 days<sup>-1</sup> | [19] |
| Sexual reproduction |  |  |  |
| $p_{inh}$ | Probability of a sexual propagule inheriting the genotype at locus $g$ of parent $Par_1$ genotype | 0.5 | Fixed |
| Pathogen dispersal |  |  |  |
| $g(\cdot)$ | Dispersal kernel | Power-law function | See note S2 |
| $\mu_{mean}$ | Mean dispersal distance | 20 m | [19] |
| $a$ | Scale parameter | 5 | Fixed |
| $b$ | Width of the tail | 3.5 | [26–28] |
| Contamination of healthy hosts |  |  |  |
| $\pi(\cdot)$ | Contamination function | Sigmoid | See note S2 |
| $\sigma$ | Parameter 1 related to the position of the inflection point | 3 | [19] |
| $\kappa$ | Parameter 2 related to the position of the inflection point | 5.33 | [19] |
| Host-pathogen genetic interaction |  |  |  |
| $G$ | Total number of major genes | 2 | Fixed |
| $\tau$ | Mutation probability | [10<sup>-5</sup>; 10<sup>-4</sup>]<sup>b</sup> | Varied |
| $\theta$ | Fitness cost of infectivity | [0; 0.25] | Varied |
| Fungicide treatment |  |  |  |
| $t^*$ | Application dates | {1 to 109 by step of 3 } | Fixed |
| $\epsilon_{eff}$ | Fungicide maximal efficiency | 1.0 | Fixed |
| $\epsilon_d$ | Fungicide degradation rate | 0.004 | Fixed |
| $\epsilon_t$ | Fungicide application threshold | 0.05 | Fixed |
| $a$ | Parameter 1 related to the shape of the logistic function | 4.0 | [29] |
| $b$ | Parameter 2 related to the shape of the logistic function | 8.5 | [29] |

<sup>a</sup> Depending on the landscape considered (“Central”, “Diversified”, Square). <sup>b</sup> Depending on the landscape considered (“Central”, “Diversified”, Square). <sup>b</sup> Varied only in the square landscape. In Buzet landscapes (“Central”, “Diversified”), only the value 10<sup>-5</sup> was used.

#### 3.3.4 Model parameterization for the real landscapes of Buzet and deployment strategies

For the simulation of epidemics in a heterogeneous landscape, *landsepi* requires the spatial coordinates of the fields composing the landscape and information about the allocation of cultivars in these fields. For example, in a previous study [19] built-in landscapes were used in which the number, area and shape of fields were generated with a T-tessellation algorithm [30]. In another study [15], simpler landscapes consisting of identical square fields were used. By contrast, we used the coordinates of the polygons outlining the boundaries of each field to incorporate two real landscapes from the Buzet area (Fig. 1A). In a complementary manner, *landsepi* also requires a matrix defining the allocation of cultivars to the different fields for each year of simulation. These two elements (maps of the landscape and a crop-type allocation matrix) provide a representation of finely tuned deployment strategies fitting the narrative scenarios arising from discussions with the staff of the wine cooperative.

#### 3.3.5 Model outputs

We defined the evolutionary, epidemiological, environmental and economic outputs assessed at the end of a simulation run. The *landsepi* model is stochastic. We therefore performed 50 replicates *r* = 1..50 for each parameter combination. We then defined the summary statistics used to characterize the outputs over the 50 replicates. Note that *r* indices are omitted in some parts of the next section to avoid overloading notations. Similarly, as a single cultivar was assigned to each field *i* in a given year *y* in the scenarios considered, the cultivar index *v*(*i, y*) = {*SC*; *RC*} is omitted from the following equations whenever possible.

For evolutionary output, we studied the establishment of SP by defining *E*_*SP*_, a binary variable set to 1 if the SP becomes established before the end of a simulation run and 0 otherwise. The SP is considered to be established if the number of resistant hosts infected by SP exceeds a threshold (i.e. 50,000 infections, as defined by [19]) above which extinction within a constant and infinite host population becomes unlikely. In the results, we present the probability of SP establishment *p*(*E*_*SP*_) estimated as the mean of *E*_*SP*_ over the 50 replicates performed.

For the epidemiological output, we used the area under the disease progress curve (AUDPC) as a measurement of disease severity. The AUDPC represents the average (over the cropping seasons) proportion of the existing hosts displaying disease [30]. For each replicate, we assessed the percent change in AUDPC relative to the mean under the baseline scenario *S*0 as 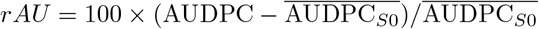 where 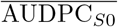 is the mean AUDPC under the baseline scenario *S*0 over 50 replicates.

For the environmental output, we assessed the treatment frequency index (TFI), estimated as the annual number of fungicide applications/field, averaged over the whole landscape and all cropping seasons. We then assessed the percent change in TFI relative to its mean under the baseline scenario *S*0 as 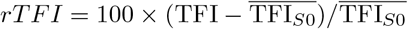 where 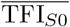 is the mean TFI under *S*0 over 50 replicates.

For the economic output, we evaluated cumulative discounted net returns over the years studied. Annual net revenue depends on the yield (tonnes), which we computed for each field *i* and year *y* as:

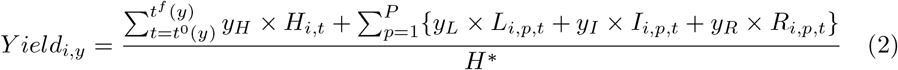

Where *y*_*H*_, *y*_*L*_, *y*_*I*_ and *y*_*R*_ are the theoretical yields of cultivar *v*(*i, y*), grown in field *i* in year *y*, associated with health statuses H, L, I and R, respectively. *H*^∗^ (ha^-1^), is the cumulative number of hosts/ha for a season in the absence of disease, set to 1780.9 × 10^4^ (Supporting Information 6). The terms *t*^0^(*y*) and *t*^*f*^ (*y*) denote the first and last days of the cropping season *y*, respectively.

Annual net revenue NR (€) was then calculated for each field *i* and year *y* as:

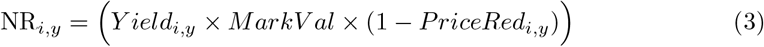

Where *MarkV al* is the market value per tonne of yield and *PriceRed*_*i,y*_ is the discount rate applied to the price paid to the grower if the disease severity on clusters at harvest exceeds a particular threshold. This made it possible to account for the yield quality loss caused by GDM [31]. The calculation of *PriceRed*_*i,y*_ is detailed in the Supporting Information 6.

We then calculated the costs (€) for each field *i* and year *y* by summing the costs due to the application of fungicide treatments (*OperationalCost*) and the cost for uprooting and replanting the cultivar (*PlantingCost*):

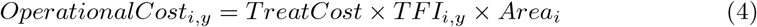

Where *TreatCost* is the cost (€/ha) for a single application of fungicide treatment, *TFI*_*i,y*_ is the number of treatments applied in year *y* and field *i* and *Area*_*i*_ is the area (ha) of field *i*.

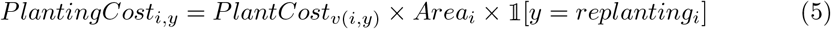

Where *PlantingCost*_*v*(*i,y*)_ is the cost (€/ha) of uprooting and replanting cultivar *v* in field *i* and year *y*, and 𝟙[*y* = *replanting*_*i*_] is an indicator function, taking a value 1 if year *y* corresponds to the year in which the vines in the field *i* are uprooted and replanted, and 0 otherwise.

Finally, we calculated the cumulative discounted stream of net returns or simply cumulative net benefits (NB_*i*_) for the field *i* (€) as previously described [32], as:

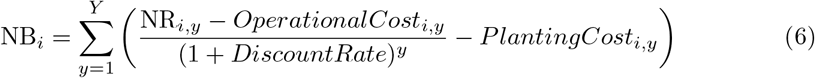

Where *DiscountRate* is the real discount rate, corresponding to the interest rate used to calculate the current value of future cash flows from a project or investment. Note that, in this equation, the planting costs are considered as a long-term capital investment whereas operational costs are considered as short-term recurring expenses. The discount rate allows operational costs to be treated as investments. Finally, the NB calculated for each field *i* were aggregated at the the scale of the cooperative (€):

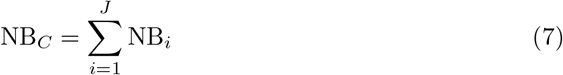

We assessed the percent change in NB relative to its mean value under the baseline scenario *S*0 as 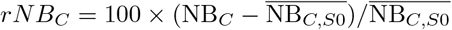 where 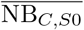 is the mean net benefit under S0 over 50 replicates.

Finally, we calculated the net benefit NB_*f*_ for each farmer *f* by applying equation 7 to the subset of fields *i* belonging to farmer *f* = 1..57 in the “Central” vineyard or *f* = 1..36 in the “Diversified”) vineyard and then the percent change in NB relative to the mean under S0 as 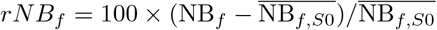 where 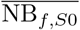 is the mean NB for farmer *f* under the baseline scenario *S*0 over 50 replicates.

The parameters used to calculate economic output are presented in Table 4. We assumed that the only difference between RC and SC was the planting cost *PlantCost*_*v*_, which is 2.5 times higher for RC [33]. In the multi-criteria performances reported for the scenarios tested, the values reported for the epidemiological, environmental and economic outputs are medians (across 50 replicate simulations), unless otherwise indicated.

**Table 4.** Summary of the parameters used to calculate economic output

| Notation | Parameter | Cultivar |  | Source |
| --- | --- | --- | --- | --- |
|  |  | SC | RC |  |
| $y_H$ | Theoretical yield for H host (tonnes/ha) | 6.7<sup>a</sup> | 6.7 | Buzet |
| $y_L$ | Theoretical yield for L host (tonnes/ha) | 6.7 | 6.7 | Buzet |
| $y_I$ | Theoretical yield for I host (tonnes/ha) | 6.7 | 6.7 | Buzet |
| $y_R$ | Theoretical yield for R host (tonnes/ha) | 0 | 0 | Buzet |
| $PlantCost_v$ | Uprooting and replanting cost (€/ha) | 5481<sup>b</sup> | 13,703<sup>b</sup> | Buzet |
| $MarkVal$ | Market value of yield (€/tonne) | 600 | 600 | [31] |
| $TreatCost$ | Cost of fungicide treatment (€/treatment/ha) | 67 | 67 | Buzet |
| $DiscountRate$ | Discount rate | 0.04 | 0.04 | [34] |

<sup>a</sup> The theoretical yield (tonnes/ha) is obtained by the division $6.7=50/7.5$ where 50 is the mean yield of wine (hl/ha) and 7.5 is the volume (hl) of wine obtained from 1 tonne of grapes.
<sup>b</sup> The planting cost differs between cultivars $v = \{SC; RC\}$ due to the price of plants in the nursery. For SC, the planting cost is $5481 = 4350 \text{ plants/ha} \times 1.26 \text{ €/plant}$ and for RC it is $13703 = 4350 \text{ plants/ha} \times 1.26 \text{ €/plant} \times 2.5$ .

### 3.4 Effects of fungicide and the massive or progressive introduction of RC

Our results for the scenarios studied in the Buzet wine cooperative landscape led us to study more specifically the effects of two factors on resistance durability and disease control by running simulations on a level playing field: (i) presence or absence of fungicide application based on a severity threshold and (ii) the massive or progressive introduction of RC for strategies characterized by the same mean proportion of RC cultivated over 30 years. These comparisons were conducted in a theoretical agricultural landscape of 225 ha (1500 × 1500 m) consisting of uniform square fields with an area of 0.5625 ha (75 × 75 m). This landscape consisted of 400 fields. The model was used to assess the evolutionary and epidemiological outputs (see above) for 336 parameter combinations resulting from a full factorial design combining (i) presence or absence of fungicide use (2 levels), (ii) fitness costs (3 levels: 0, 0.05, 0.25), (iii) probability of mutation (2 levels), (iv) the massive or progressive introduction of RC (2 levels) and (v) the mean percentage of the landscape under RC over a period of 30 years (14 levels). The corresponding dynamics of RC introduction into the landscape are illustrated in figure 2. We performed 120 replications for each parameter combination to estimate the probability of SP establishment (*p*(*E*_*SP*_)), the mean time to establishment given that the SP becomes established 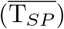 and the mean percent change in AUDPC relative to the mean under the baseline scenario *S*0 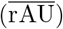. We also compared the occurrence of SP establishment with and without fungicide treatment, based on relative risks. Relative risks are the ratio of the probability of an outcome of interest (here SP establishment) in an exposed group (here in the presence of fungicide treatments) to the probability of this outcome in the control group (here in the absence of fungicide treatments). We present below the mean relative risks averaged over the type of introduction and the cropping ratio considered. We also calculated the relative risks of of SP establishment with a progressive (exposed group) and a massive (control group) strategy of RC introduction, and we present the mean relative risks by treatment and cropping ratio.

**Fig 2.**
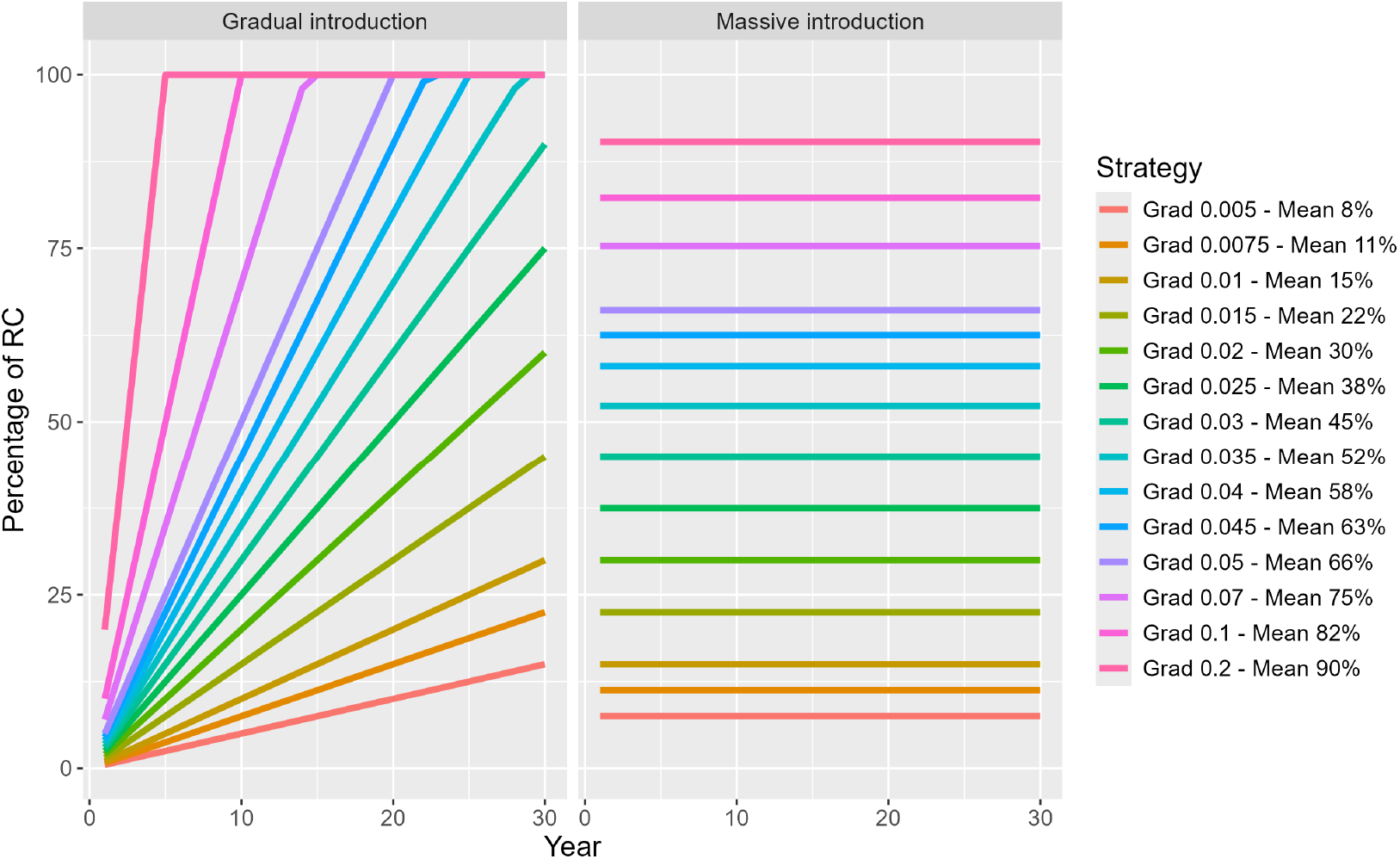
Dynamics of RC introduction into the landscape according to a progressive (left panel) or massive (right panel) strategy for 14 mean percentages of RC in the landscape over a period of 30 years. The mean percent RC for the progressive (mean over 30 years) and massive strategies is indicated in the legend after the “Mean”. The numerical value after “Grad” corresponds to the proportion of fields planted with RC each year in the progressive strategy.

## 4 Results

### 4.1 Scenarios for resistant cultivar deployment defined for Buzet

The scenarios to be tested were developed on the basis of discussions with the team members of the cooperative during the second and third workshops. Right from the start of the second workshop, discussions focused on the possible options for the deployment of a RC pyramiding the resistance genes *Rpv1* and *Rpv3*.*1*, the only cultivar for which the cooperative requested PDO authorization. Six deployment scenarios were retained after the third workshop (Table 5), reflecting strategies of particular interest to the cooperative team. These scenarios were based on (i) the age of the vines in the targeted fields, (ii) whether or not PDO rules applied, (iii) the replanting rate, or (iv) the specific location of the field. The scenarios were defined based on a “plantation rule” guiding the choice of fields in which the RC would be planted in each of the years of the simulation *y* = 1..30. Below, fields containing vines more than 30 years old (*i*.*e*. planted more than 30 years before a given year *y*) are referred to as “fields ≥ 30 years”. All the scenarios begun with only SC in the landscape (year 0), corresponding to the current situation.

**Table 5.** Summary of the scenarios coconstructed with the team from the wine cooperative. Scenarios are defined based on a planting rule guiding the choice of fields in which RC or SC are planted. We refer to fields that were planted more than ≥ 30 years before a given year of simulation *y* as “fields ≥ 30 years”. The total percentage of the area of the territory planted with RC at year 30 is reported, along with the mean percentage of the area of the territory planted with RC over each of the 30 years of simulation. In addition, the total number of fields under RC at year 30 is reported. The scenarios concern the “Central” (1422 fields) and “Diversified” (709 fields) vineyards.

| Name Plantation rule |  | Central vineyard |  |  | Diversified vineyard |  |  |
| --- | --- | --- | --- | --- | --- | --- | --- |
| | | %RC at<br>$y = 30$ | mean<br>%RC | RC plots<br>at $y = 30$ | %RC at<br>$y = 30$ | mean<br>%RC | RC plots<br>at $y = 30$ |
| S0 | SC planted in fields $\geq 30$ years | 0 | 0 | 0 | 0 | 0 | 0 |
| S1 | RC planted in fields $\geq 30$ years | 100 | 80 | 1422 | 100 | 88 | 709 |
| S2 | RC planted in 3.3%/year of the oldest fields | 98 | 50 | 1410 | 96 | 54 | 690 |
| S3 | RC planted in fields $\geq 30$ years up to 5% of the area of each farm, otherwise SC planted | 5 | 4 | 237 | 4 | 4 | 107 |
| S4 | RC planted in fields $\geq 30$ years up to 20% of the area of each farm, otherwise SC planted | 19 | 19 | 575 | 19 | 19 | 273 |
| S5 | RC planted at year 1 in fields classified as NTZ Aqua<sup>a</sup>, SC planted in fields $\geq 30$ years | 9 | 9 | 124 | 5 | 5 | 21 |
| S6 | RC planted at year 1 in fields classified as NTZ Riverain<sup>b</sup>, SC planted in fields $\geq 30$ years | 6 | 6 | 76 | 4 | 4 | 27 |

<sup>a</sup> NTZ Aqua: fields with restrictions on treatments due to proximity to water sources; <sup>b</sup> NTZ Riverain: fields with restrictions on treatments due to proximity to residential areas (*e.g.* houses, schools).

In scenario S1, year of simulation *y*, all fields ≥ 30 years were planted with RC. In scenario S2, replanting occurred every year in 3.3% of the oldest plots, ensuring a complete renewal of the plots over a period of 30 years. In scenarios S3 and S4, year of simulation *y*, all fields ≥ 30 years were planted with RC until proportions of 5% and 20% of the vineyard area for each farm, respectively, were attained. Once the thresholds (5% or 20%) were reached, the fields ≥ 30 years were replanted with SC during the remaining years of the simulation. In scenarios S5 and S6, RC were planted in year 1 in fields located in the no-treatment zone (NTZ) in which fungicide use is restricted due to proximity to watercourses or elements of the hydrographic network (“NTZ Aqua”) or human dwellings (“NTZ Neighbor”), regardless of the age of the vines planted in the fields. Conversely, for subsequent year of simulation *y*, all the remaining fields ≥ 30 years were replanted with SC. In line with current regulations, we assumed that copper fungicides (authorized for use in both conventional and organic farming) could be used in the NTZ, as only synthetic pesticides are prohibited in the NTZ. Finally, we simulated a baseline *status quo* scenario S0 in which only SC were deployed. In S0, year of simulation *y*, fields ≥ 30 years were replanted with SC. This scenario corresponds to typical current practices for vineyard renewal in this cooperative and provides a reference point for comparison with the results obtained for the other scenarios.

The scenarios differed considerably in RC planting dynamics over the 30-year period (Fig. S2). Four scenarios (S3, S4, S5, S6) were characterized by less than 20% of the total area being under RC by year 30, whereas more than 96% of the total area was under RC by this time point in scenarios S1 and S2. Only scenario S2 was characterized by a progressive introduction of the RC. The other scenarios involved a rapid introduction of RC right from the start of the simulation due to the age distribution of the vines in the fields. In particular 65% of the area of the “Central” vineyard was planted with RC after year 1 with scenario S1 but only 15% with scenario S2. The scenarios also differed in terms of the replantation of SC. Fields ≥ 30 years were replanted with SC in scenarios S3, S4, S5, S6 and S0, whereas no such replantation occurred in S1 and S2.

The scenarios also differed in terms of specific features of the landscape configuration. In particular, the fields located in the NTZ were fewer in number but larger than those in other areas. Accordingly, although scenarios S3, S5 and S6 had similar areas under RC, the number of fields planted with RC in S3 was greater (and these fields had a smaller mean size) than in scenarios S5 and S6 (Table 5). These differences translate into differences in the probability of pathogen propagule dispersion from fields cultivated with SC to fields cultivated with RC (Table S1).

### 4.2 Multicriteria performances of the six scenarios

#### Durability of the resistant cultivar

The probability of SP establishment varied considerably between scenarios and fitness costs, with a lesser influence of the subdomain of the vineyard considered (“Central” or “Diversified” vineyard) (Fig. 3, top row). In the “Central” vineyard, the probability of SP establishment ranged from 0.02 (scenario S6, high fitness cost) to 0.72 (scenario S4, intermediate fitness cost). The effect of fitness cost was generally consistent between scenarios: the higher the fitness cost, the lower the probability of SP establishment. Conversely, considerable variation for the probability of SP establishment was observed between scenarios for a given fitness cost, for example from 0.24 to 0.62 for a fitness cost of zero and from 0.02 to 0.32 for a high fitness cost. These variations were not correlated with the mean percent of the territory surface planted with RC over the 30 years of the simulation or with the maximum percentage. Finally, the probability of SP establishment generally tended to be lower in the “Diversified” than in the “Central” vineyard. The mean probability of SP establishment for the six scenarios and the three fitness costs in the “Central” (0.28) vineyard was twice that in the “Diversified” (0.14) vineyard.

**Fig 3.**
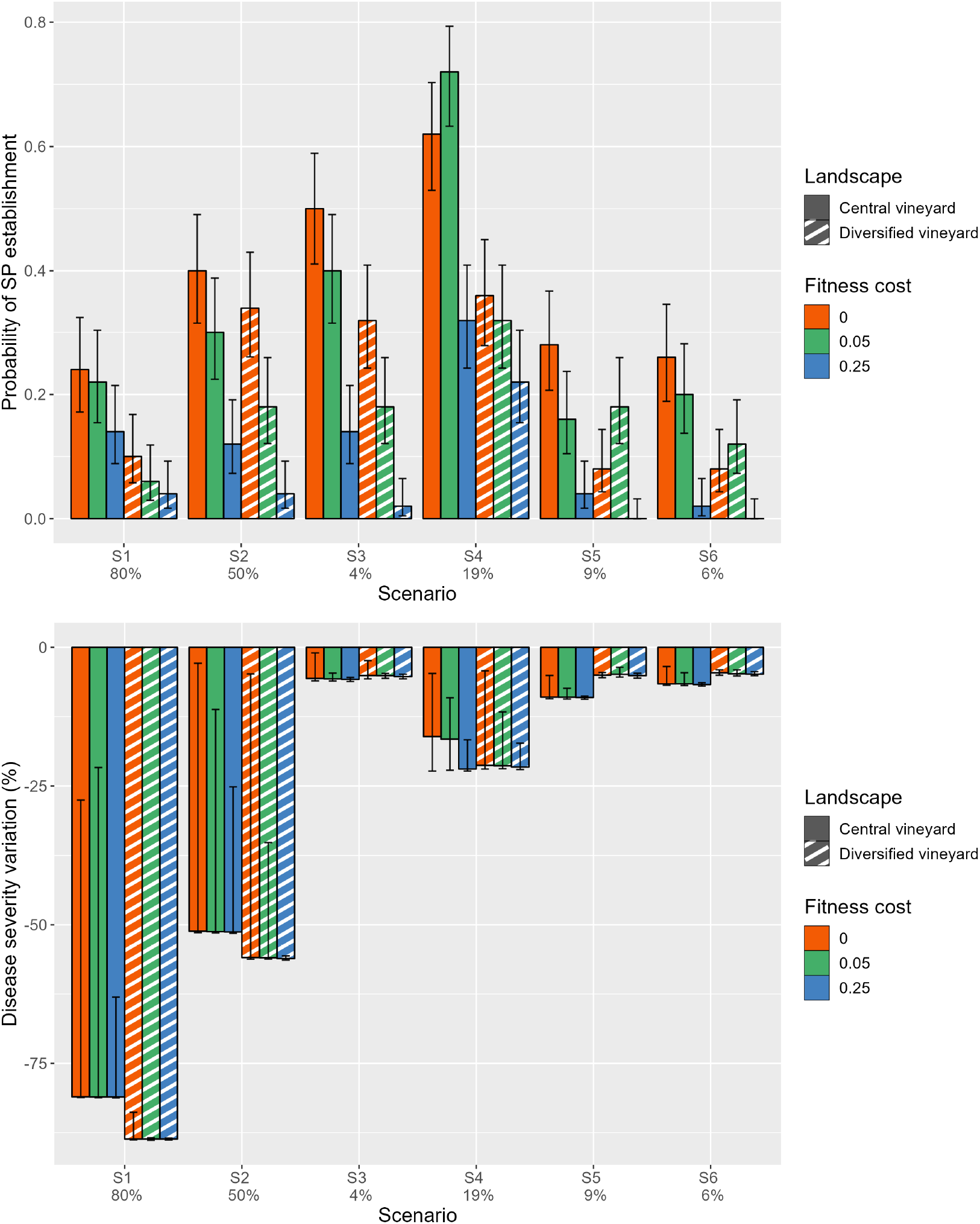
Evolutionary (top row) and epidemiological (bottom row) outputs for the six scenarios tested. The evolutionary output considered is the probability of SP establishment (*p*(*E*_*SP*_)). The epidemiological output considered is the percent change in the area under the disease progression curve relative to the baseline strategy S0 (*rAU*). Simulations were performed in the two landscapes of the cooperative territory and for three fitness costs. Barplots indicate the median of 50 replicates along with the 10% and 90% quantiles. The mean percentage of the cooperative territory planted with RC over the 30 years considered in the “Central vineyard” landscape is indicated on the x-axis.

#### Epidemiological control

In the baseline scenario S0, a low-intensity GDM epidemic is developing, and fungicide treatment are applied to all SC fields to keep the proportion of infected hosts below 5%. Similarly, SC and RC fields are treated in scenarios S1 to S6 as soon as this 5% infection threshold is reached. Therefore, any differences in the epidemiological, environmental, or economic outcomes in scenarios S1 to S6 result from the effect of disease resistance, because treatment practices were uniform across cultivars. Relative to S0, the decrease in disease severity obtained ranged from 5% to 89%, indicating that deploying RC according to any of the scenarios considered improved epidemiological control (Fig. 3, bottom row). The median epidemiological control was only marginally affected by the landscape considered. Scenarios S1 and S2 provided the best epidemiological control, with decreases of 81% to 89% depending on the landscape for S1 and of 51% to 56% with S2. By contrast, scenarios S3 to S6 provided much lower levels of epidemiological control, with the decrease in severity ranging from 5% to 22%. This finding is consistent with the lower proportion of RC planted in these scenarios.

The variability of disease control across replicates, as illustrated by the 80% confidence interval (Figure 3, bottom row), decreased with fitness cost and was much greater in scenarios S1 and S2. In these two scenarios, in which the pattern was easy to observe, the 50 simulation replicates could be split into two groups one with and the other without resistance breakdown. Disease severity was very low in the set of simulations without breakdown, as most fields were planted with RC in these scenarios. Conversely, the benefit of the RC was lost across large portions of the landscape in the set of replicates with breakdowns, in which disease severity increased with the speed of resistance breakdown, with rapid breakdowns resulting in disease levels similar to those in the S0 scenario. The resulting bimodal distribution of disease severity (Fig. S3) accounts for the large confidence intervals obtained with S1 and S2. Moreover, the variability of disease control decreased with increasing fitness cost, because the probability of SP establishment — and consequently the number of replicates displaying resistance breakdown — tended to decline with higher fitness costs. By contrast, the median level of epidemiological control remained unaffected by fitness cost as fungicides were applied on the RC in cases of breakdown.

#### Fungicide applications

The mean number of fungicide applications/field/season for the baseline scenario S0 was 10.1 and 9.9 for the “Central” and “Diversified” vineyards, respectively. These values are consistent with the number treatments applied over French vineyards in 2010 (TFI = 10.1 ± 5, [5]). RC deployment decreased the number of fungicide applications, as expected (Fig. 4, top row). The largest decreases in fungicide application, of 83% and 50%, were observed for scenarios S1 and S2 and correspond to 1.7 and 5.0 applications/field/season, respectively, in the “Central” vineyard. The decrease in fungicide use was closely linked to the level of disease control achieved (Pearson correlation coefficient = 0.98), as fungicide was applied based on a disease severity threshold rather than on a fixed schedule. The large confidence intervals for the percent change in the number of fungicide applications for scenarios S1 and S2 have the same underlying cause as those for disease severity: the occurrence or absence of resistance breakdown (Fig. S3).

**Fig 4.**
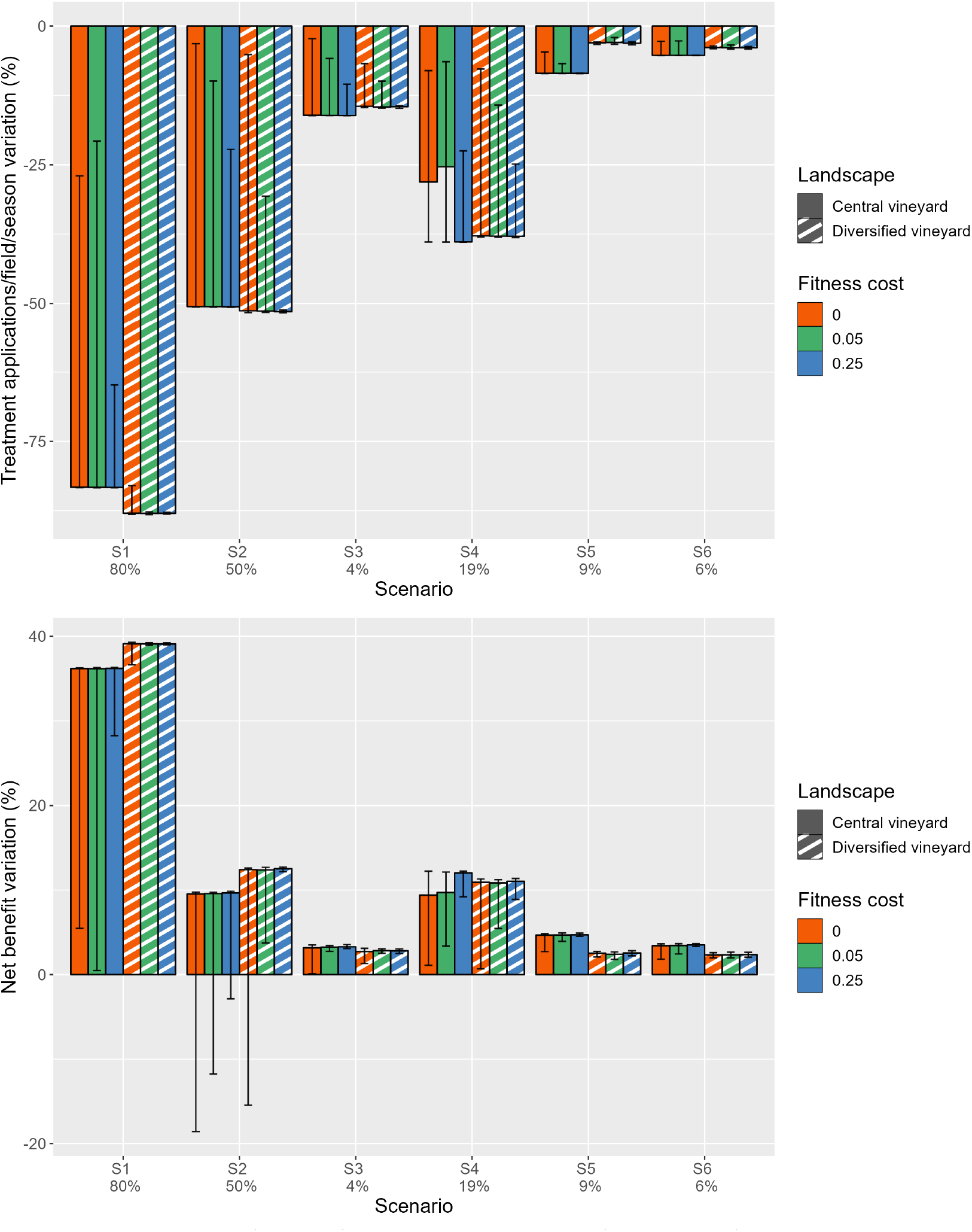
Environmental (top row) and economic output (bottom row) for the six scenarios tested at the scale of the cooperative. The environmental output considered is the percent change of the mean number of treatment applications/field/season relative to the baseline strategy S0 (*rTFI*). The economic output considered is the percent change in net benefit relative to the baseline scenario S0 at the scale of the cooperative (*rNB*_*C*_). Simulations were performed in the two landscapes of the cooperative territory (“Central” and “Diversified” vineyards) and for three fitness costs. Barplots indicate the median of 50 replicates along with the 10% and 90% quantiles. The mean percentage of the area of the cooperative territory planted with RC over the 30 years considered in the “Central” vineyard landscape is indicated on the x-axis.

#### Net benefits

The deployment of RC increased the net benefits at cooperative scale relative to the baseline scenario S0 (Fig. 4, bottom row). Scenario S1 provided the largest increase in median net benefits (36% with respect to S0 in the “Central” vineyard). This value fell to 13% for scenario S2. As described above, these two scenarios were also characterized by a large variability between replicates, especially for null and intermediate fitness costs. In particular, in the “Central” vineyard, scenario S2 resulted in financial losses in 36% of the 50 replicates for fitness costs of zero and 28% of the 50 replicates for a fitness cost of 0.05, indicating that RC deployment leads to higher costs than maintaining the status quo. The higher probability of SP establishment in scenario S2 than in scenario S1 and the progressive introduction of RC in S2 increase the likelihood of overlapping costs: the cost of fungicide applications for SC and for RC in cases of resistance breakdown,together with 2.5 times higher planting costs for RC. When combined, these costs may exceed those incurred in S0, particularly if the resistance is rapidly broken down. By contrast, the percent change in net benefits in scenario S1 was generally positive, with less than 10% of the replicates leading to financial losses. The massive initial introduction of RC in S1 strongly decreased the cost of fungicide applications, a saving that mostly more than compensated for the higher costs of planting RC. The increase in median net benefit in scenario S4 was similar to that in S2 with a lower proportion of RC planted (19% of RC in S4 and 50% in S2; Table 5). Moreover, less than 8% of replicates in S4 (assuming no fitness cost) resulted in financial losses, versus 36% in S2. The increase in median net benefit provided by S3, S5 and S6 was below 5%. These relatively weak economic performances can be attributed to the limited proportion of the landscape allocated to RC in these scenarios. Finally, even though fitness cost did not generally affect median net benefit, higher fitness costs reduced output variability and, in particular, the frequency of financial losses.

A different picture emerged at farm scale when we considered the percent change in median net benefits (over 50 replicates) calculated for each farm. Most farms benefited from the introduction of RC, but some farms experienced losses in median net benefit relative to the baseline scenario (Fig. S4). For instance, in scenario S2, up to 22 of the 57 farms in the “Central” vineyard experienced decreases in median net benefits. This was the case for 13 of the 36 farms in the “Diversified” vineyard.

### 4.3 Effects of fungicide treatment and of the massive or progressive introduction of RC

The scenarios proposed by the team of the Buzet wine cooperative led us to study more specifically the effects of (i) the presence or absence of fungicide applications (based on a severity threshold) and (ii) the massive or progressive introduction of RC. These comparisons were performed for evolutionary and epidemiological outputs on a level playing field for deployment strategies characterized by the same mean percentage of the total area under RC over a period of 30 years (hereafter referred to as the cropping ratio). Below, we focus particularly on the results obtained with the intermediate probability of mutation (10^-5^, the same value as used previously).

The relationship between cropping ratio and the probability of SP establishment yields a bell-shaped curve, with peak probabilities observed at intermediate cropping ratios for all combinations of fitness costs, strategies of RC introduction and pesticide application tested (Figure 5 first row). The probabilities of SP establishment were generally lower when fungicide treatments were applied. The mean relative risk (averaged over the type of introduction and the cropping ratio considered) amounted to 0.70 (null fitness cost), 0.73 (intermediate fitness cost) or 0.62 (high fitness cost), suggesting a 27% to 38% decrease in the risk of SP establishment when fungicides were applied. These differences tended to disappear with the progressive introduction of the RC with a cropping ratio ≥ 80%. The probability of SP establishment was also generally lower for the progressive introduction of RC. The mean relative risk (averaged for the presence or absence of treatment and the cropping ratio considered) was 0.81 (null fitness cost), 0.79 (intermediate fitness cost) or 0.69 (high fitness cost), suggesting a 19% to 31% decrease in the risk of SP establishment for a progressive introduction. These differences tended to disappear for a cropping ratio ≤ 20%.

**Fig 5.**
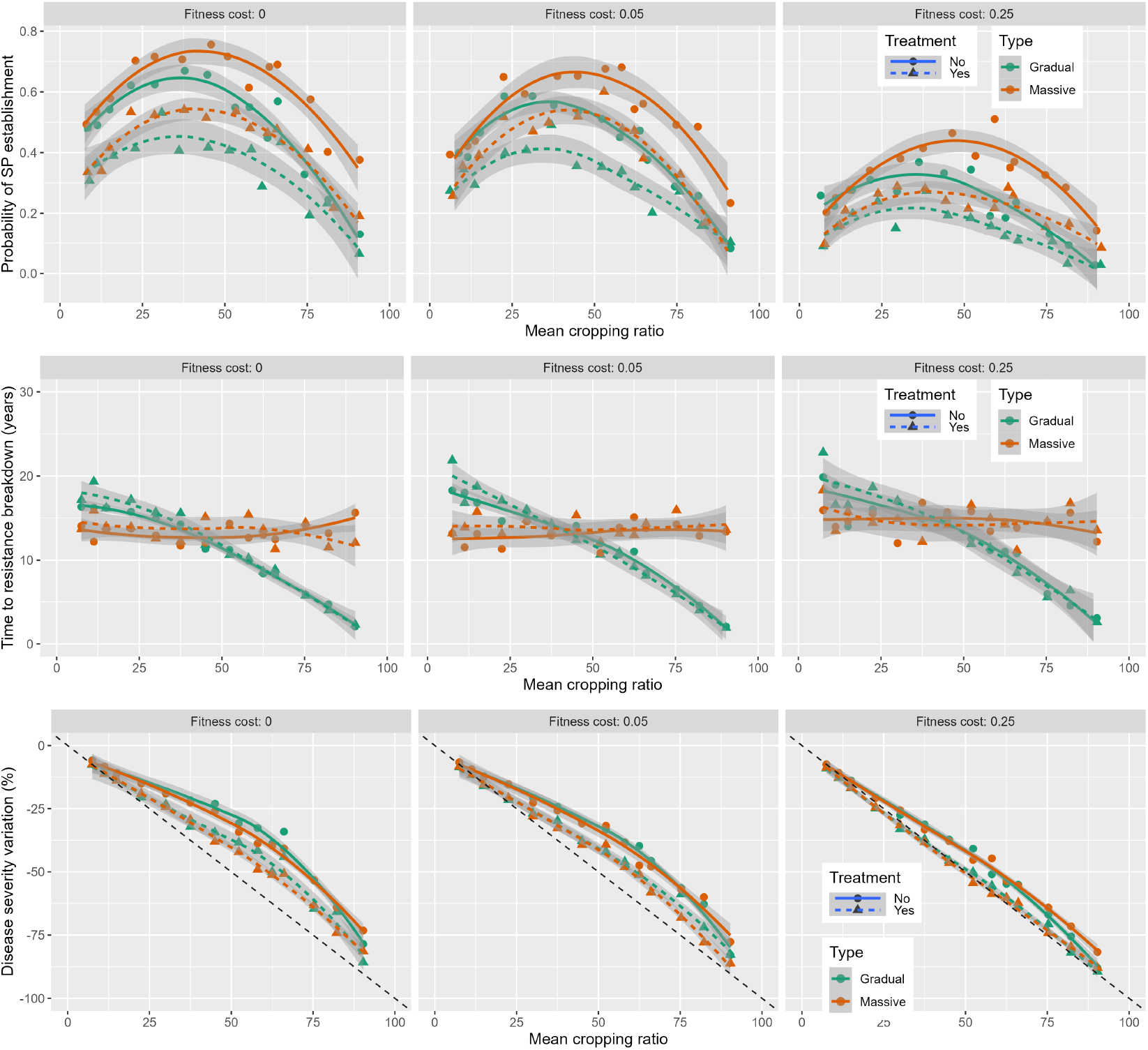
Evolutionary and epidemiological performance of deployment strategies combining massive or progressive RC introduction with the presence or absence of fungicide treatments at an intermediate probability of mutation. Probability of SP establishment (first row), time to SP establishment given that establishment occurs (second line) and percentage change of the area under disease progress curve compared to the baseline strategy S0 (third row) for three fitness costs as a function of the mean percentage of RC in the landscape over a period of 30 years. The curves are based on a locally weighted smoothing (loess) method fitted to *p*(*E*_*SP*_), 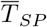 and 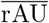 (represented by points). The shaded region represents the 95% confidence interval around the fitted values. The bisector in the third row indicates the expected variation of disease severity following the deployment of a RC with a perfect resistance impossible to break down.

The most notable difference between the progressive and massive RC introduction strategies in terms of SP establishment was the mean time to establishment in situations in which establishment occurred. The mean time to establishment, centered around 14-15 years whatever the cropping ratio for massive introduction, decreased from 16-18 years to 2-3 years with increases in cropping ratio for progressive introduction. The graph showing the dynamics of the proportion of RC over the 30-year period considered (Fig. S2) provides insight into the reasons for this difference. As shown in this figure, all progressive strategies with a mean cropping ratio ≥ 50% lead to a landscape containing only RC after a given year. As SP can only emerge from an SC field in the absence of external inoculum, the time to SP establishment is inherently constrained to be shorter than the time to the year in which RC cultivars fully occupy the landscape.

Disease severity decreases with increasing cropping ratio, following a concave relationship that becomes more pronounced as resistance breakdowns become more frequent (Fig. 5 third row). Indeed, without breakdown, disease severity would decrease strictly proportionally to cropping ratio. The use of treatments and the presence of higher fitness costs bring the relationship closer to this bisector, and their combination renders the curve indistinguishable from the bisector. By contrast, RC introduction strategy had very few, if any, effects on disease control. This may be surprising at first sight given that the probability of SP establishment was mostly higher for the massive introduction strategy. However, other processes contribute to overall epidemic intensity. First, all progressive strategies with a mean cropping ratio ≥ 50% will eliminate the pathogen in the absence of breakdown, but the benefit of the RC is lost in a larger number of years with the progressive strategy than with the massive introduction strategy for a cropping ratio ≥ 50% if resistance breakdown occurs. Furthermore, before the breakdown, the introduction of the RC decreases the intensity of the epidemic in the SC through a dilution effect. This dilution effect increases with cropping ratio, so massive introduction benefits from a stronger dilution effect than progressive introduction during the first 15 years of simulation, whereas the situation is inversed after year 15. Overall, the balanced effects of these mechanisms (probability of RC breakdown, conditional time to breakdown, dilution effect until breakdown) result in similar levels of epidemiological control for the massive and progressive strategies of RC introduction.

Finally, we tested the effect of a 10 times higher mutation probability (10^-4^). As highlighted in our previous studies [15, 19], the order of magnitude of the probability of mutation relative to pathogen population size is a key factor shaping resistance durability and disease control. In our case study, a mutation probability 10 times higher eliminated the advantage of deploying RC in most of the tested production situations, mostly because it led to a very rapid breakdown of resistance (Fig. S5).

## 5 Discussion

In this discussion, we will explore how an action-research approach was used to develop bottom-up strategies for deploying RC while encouraging researchers to enhance their model with new features. These enhancements, in turn, enabled us to address novel questions. Specifically, we examined the effects of (i) a massive or progressive introduction of RC and (ii) fungicide treatments on extending resistance durability. The results for the strategies tested in the Buzet vineyard will not be discussed in detail, as they are highly specific to this case study.

Our aim was to compare bottom-up strategies for deploying RC identified during an action-research approach grouping together a farmer’s organization and researchers. We used mathematical modeling as a tool for exploring and understanding complex issues in a real agricultural landscape [17]. The modeling approach used here is based on solid mathematical foundations and a knowledge of the epidemiology and evolution of plant pathogens acquired over the last four decades [13]. Modeling is free from the logistic, financial and legal constraints associated with experimentation in agricultural systems and can therefore be used to compare deployment strategies without the risk of disastrous epidemics or resistance breakdowns. An action-research involving a partnership with key local stakeholders was previously used to build scenarios combining RC, cropping practices and spatial territory organization for managing *Phoma* stem canker on oilseed rape [35]. A dedicated simulator was used during the workshops [36, 37]. In the Netherlands, workshops with both conventional and organic farmers explored strategies for deploying late blight-resistant potato cultivars [38, 39]. In our case study, the ability to perform real-time simulations on the actual plots of Buzet facilitated dialog with the staff of the wine cooperative. The involvement of an agency specializing in participatory workshops enabled each participant to focus on the core aspects of their profession, thereby addressing a common challenge in action-research, in which experts often lack the social experience, interaction skills, or natural ability to collaborate effectively with local participants [40].

The first workshop highlighted the need to upgrade *landsepi* with two features to increase its acceptance among stakeholders. First, the integration of fungicide treatments was necessary to allow correct comparisons to be made with the reference scenario, in which about 10 treatments are typically used to control GDM. Fungicides, which remain an important component of integrated pest management in vineyards [29], are rarely considered in studies on resistance deployments despite their potential to enhance resistance durability [13]. Chemical applications, especially on fields planted with SC, increases the durability of both qualitative and quantitative resistances [41, 42]. Our results are consistent with these previous findings. The mean (over cropping ratio and type of introduction) relative risk of SP establishment decreased by 27% to 38% (depending on fitness costs) when fungicides were used (Fig. 5 first row). However, the changes in these values with changes in the threshold for fungicides application (*ϵ*_*t*_) have yet to be studied, together with the extent to which lowering this threshold could offset low fitness costs. Similarly, the modeling of treatments might reflect current options more accurately (e.g. fungicides with post-infection activity, systemic fungicides) [29, 43], as well as allowing different treatment programs for different cultivars. One previous study [38] found that fields planted with potato cultivars resistant to late blight were not infected when fungicides were applied to all susceptible fields because the disease was successfully suppressed, resulting in a very small virulent population that was unable to infect the fields planted with resistant varieties, or if fungicide was sprayed on all RC fields because this would prevent the infection of the RC and the spread of the virulent strain. Future studies could test strategies in which RC are specifically treated later in the fall with fungicides with post-symptom activity to reduce the size of the GDM population just before the sexual reproduction phase.

Second, the first workshop also suggested that the introduction of RC over cropping seasons should be progressive. This is “the natural scenario”. Most studies have considered a static landscape (i.e., cultivar allocation to the fields is fixed) [13], but some models have demonstrated that resistance durability can be improved by varying the proportion of resistance used in mixture or landscape mosaics each year [44, 45]. However, to the best of our knowledge, only one study [46, 47] has specifically investigated scenarios mimicking a progressive introduction of RC. Using a gene-for-gene model originally accounting for the coevolution of resistance and virulence in natural plant diseases, the authors showed that the addition of a percentage of resistant genotypes at each generation led to “boom and bust” cycles typical of agroecosystem. With a different modeling framework, another study [39] also showed that the progressive adoption by farmers of a potato cultivar resistant to late blight leads to “boom and bust” cycles: the percentage of farmers growing the RC increased until resistance breakdown occurred. Here, we compared massive and progressive introductions of RC for strategies with the same mean cropping ratio of RC over a period of 30 years. Our simulations indicated that a progressive introduction worsened evolutionary control. Lower cropping ratios of RC in the landscape and lower rates of RC introduction (*i*.*e*., a longer coexistence between SC and RC), resulted in a higher risk of virulent strains emerging from fields of SC and spreading to neighboring fields of RC. In human disease control, the design of vaccination strategies places significant emphasis on the speed of vaccination, akin to the rate of RC introduction in our case study. The risk of spread of a vaccine-adapted variant is maximal for an intermediate speed of vaccination rollout [48, 49]. This result is probably linked to the shape of the relationship we observed between the rate of RC introduction and the probability of SP establishment (Fig. 5 first row). The progressive introduction of RC results in a higher probability of resistance breakdown, but not a significant worsening of epidemiological control due to the balance between factors (*e*.*g*. timing of SP establishment, intensity of the dilution effect), sometimes favoring the progressive strategy and sometimes favoring the massive introduction strategy. In particular, the progressive strategy can eventually lead to a monoculture of RC (the strategies with mean cropping ratio *>* 0.5 in Fig. S2 left panel), potentially eliminating the pathogen before adaptation occurs. An open issue remains regarding the impact of migration introducing an external inoculum source on these results.

Third, the first workshop outlined the need to compare the strategies on the basis of economic outputs we added in the model. This need contrasts with the scarcity of models taking into account the socioeconomic context (only six of the 69 studies reviewed by [13]). The link between epidemiology and economics is traditionally established through the estimation of crop yields and harvest losses caused by diseases [50]. The most detailed studies on this aspect to date are those conducted on potato late blight in the Netherlands [38, 39, 51] and blackleg of winter oilseed rape in France [36, 37]. In these models, crop yield was calculated as a function of crop cultivar, disease severity, and climatic variables. However, data on the effects of these variables on yield are frequently unavailable. More simply, some studies assumed that RC grow more slowly [51] or have a lower yield [52] than SC. Here, we did not consider a yield gap between RC and SC as, for grapevine, yield is restricted to well below the potential biological yield by the rules of the PDO designed to favor harvest quality. We also assumed that the rules for fungicide treatment applied in the simulation would enable PDO yield targets to be met. Interestingly, with this model, the net benefits could be estimated and compared at the scales of both the wine cooperative and individual farms. This was possible because the *landsepi* model is spatially explicit and the owners of the field plots is known. In turn, it raise the question of how to distribute private costs at the individual farm level (e.g. the additional cost of planting RC) for a collective benefit at the cooperative level (e.g. better disease control and decrease in the number of fungicide treatments) for strategies leading to uneven RC planting among farmers. Two economic mechanisms could be mobilized at the cooperative level and tested in the model to compensate for the loss of income of these farms. First, price valuation [53] would require horizontal market differentiation, with RC labeled either by their variety or their reduced treatments use. It could target consumers who value environmentally friendly wines [54, 55]. Second payments for environmental services that translate non-market external values into real financial incentives for local actors providing environmental services such as reducing phytosanitary treatments [56, 57].

Last but not least, farmers are decision-makers. We considered dynamic cultivar allocation over the seasons, but the strategies were developed initially for the next 30 years. One valuable approach would involve relaxing this assumption by allowing farmers to decide each year whether to keep cultivating SC or to plant RC according to criteria assessed at the end of the previous growing season on their farms and/or in the surrounding fields. The agent-based model developed in a previous study [39] links the decisions of farmers (notably planting a RC or a SC in the next year) to disease dynamics, making it possible to explore the feedback mechanisms occurring in this socio-ecological system. Recent studies [58, 59] have integrated game theory with compartmental epidemiological models to highlight the importance of including growers’ decisions in plant epidemic models. In particular, one of these studies [59] investigated how growers’ strategic choices between resistant and tolerant crop cultivars influence disease pressure and collective yield outcomes. Another study [60] examined the determinants and barriers to farm-level adoption of RC, showing that farmers closer to final consumers are more likely to adopt RC.

These approaches offer a promising perspective: designing policy interventions favoring the durable management of RC. These interventions could, for instance, include subsidies for the planting of RC, which are currently 2.5 times more expensive than SC, or adjustment of the price paid for RC or SC based on the adoption of specific agricultural practices (*e*.*g*., not exceeding a threshold of RC on the farm). Subsidies could also be specifically targeted to farmers more at risk of suffering financial losses when planting RC. Finally, interventions could favor agricultural extension services promoting the education of farmers and cooperation [61]. Ultimately, these incentives and interventions could be optimized to ensure that the deployment strategies adopted by farmers closely align with a best-case scenario in which farmers’ choices are optimized to maximize net benefits.

If the addition of new features is important, it is also crucial not to aim for an all-encompassing model. For example, the *landsepi* model does not take into account the effects of environmental variables (particularly meteorological variables) despite their known effects on the short-term dynamics of epidemics [43]. We focus solely on the interactions between epidemiological, evolutionary and economic aspects in the long term, with the aim of comparing deployment strategies on a level playing field. We assume that weather conditions would have affected all strategies in the same way and would not, therefore, change our conclusions. This conclusion may therefore be valid only if this assumption holds true. In our view, prioritizing comprehensive uncertainty reporting is more valuable than modeling new processes in *landsepi*. It is essential to quantify and report model uncertainties comprehensively, particularly when models are designed to guide actions [62]. The *landsepi* model inherently accounts for demographic stochasticity in pathogen dynamics, including mutation occurrence, genetic drift, and seasonal bottlenecks. However, uncertainties also stem from our limited knowledge of parameters and from the structure of the model itself. We addressed parameter uncertainties through sensitivity analysis, examining key parameters such as mutation probability and fitness costs, to alert stakeholders. However, our analysis did not explore alternative model structures. This challenge should be addressed in future studies using, for example, multimodel ensemble techniques.

## 6 Acknowledgments

We would like to thank the ‘Nous, les vignerons de Buzet’ cooperative for its warm welcome and the commitment of its teams during the spring 2023 workshops (Carine Galante, Carine Magot, Sébastien Bourguignon and Pierre Philippe). We would also like to thank Think+ (Eco-innovation agency) for organizing these workshops (Vincent Collet and Hélène Lovato), Gauthier Sabria (sociological analysis) and Louise Plantin for her illustrations. We also thanks the team managing the high-performance computing platform of the BioSP unit at INRAE (https://biosp-cluster.mathnum.inrae.fr/).

## Code availability

The scripts and instructions to generate part of the results presented are openly available in the dataverse Data INRAE at https://doi.org/10.57745/9IVLH2. Only the results for the simplified squared landscape can be reproduced. Due to privacy concerns, the actual map with real fields of the Buzet area cannot be published.

## Author Contributions

Conceptualization : AAU, ASM, FF. Formal Analysis : MZ, FF. Funding Acquisition: AAU, FF. Methodology : MZ, AAU, LP, JP, FF. Software : MZ, LR, JP, JFR. Writing – Original Draft Preparation : MZ, AAU, FF. Writing – Review and Editing : MZ, AAU, ASM, LP, JP, JFR, FF

## Supporting information

### Figures

**Fig S1.**
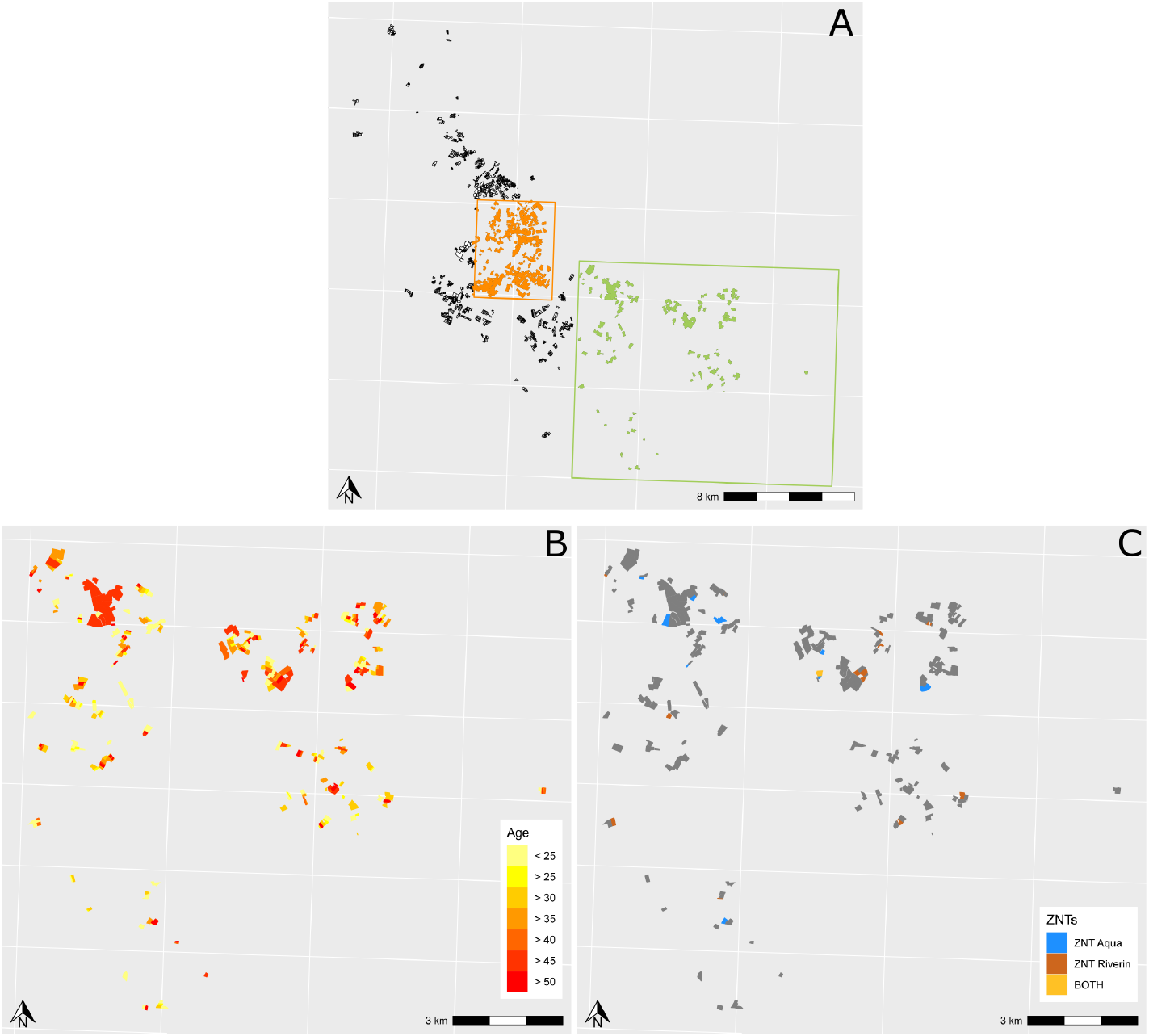
Maps of the Buzet vineyard. A: Two landscapes were considered in the Buzet territory. The “Central vineyard” corresponds to the central part of the Buzet area (in orange). This landscape (4.8 km × 6.0 km) includes 1422 fields and 57 farmers, with a mean field size of 0.43 ha. The “Diversified vineyard” corresponds to a peripheral part of the Buzet area (in green). This landscape (13.3 km × 16.0 km) includes 709 fields and 36 farmers, with a mean field size of 0.56 ha. B: Age of the vines present (“Diversified vineyard” only). C: Classification of the fields in the no-treatment zone due to proximity to an aquatic area (NTZ Aqua), human dwellings (NTZ Riverain) or both (“Diversified vineyard” only)

**Fig S2.**
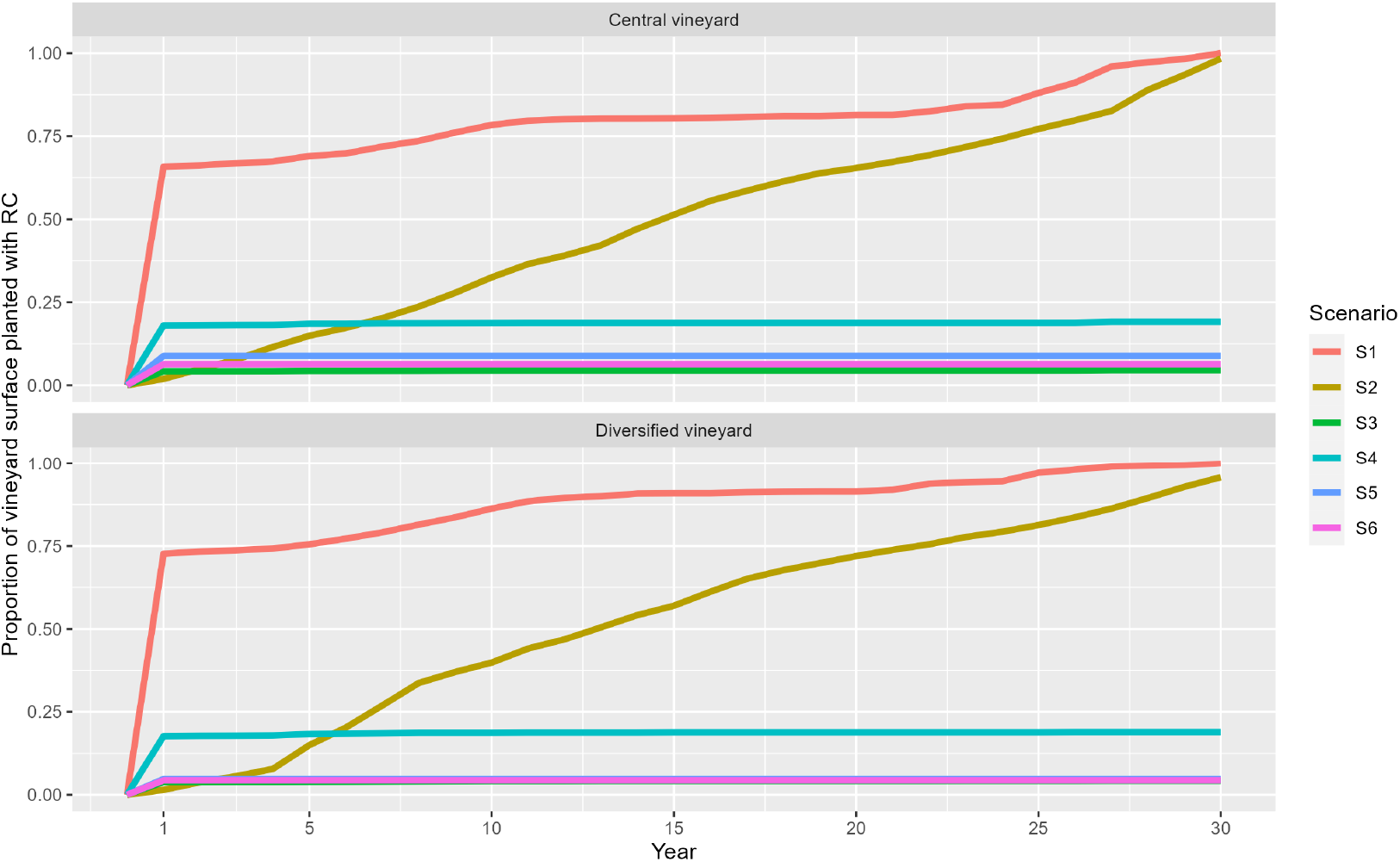
Annual dynamics of the proportion of the area under resistant cultivars across the six scenarios and the two landscapes considered. In the “Diversified vineyard”, the curves for scenarios S3, S5 and S6 overlap.

**Fig S3.**
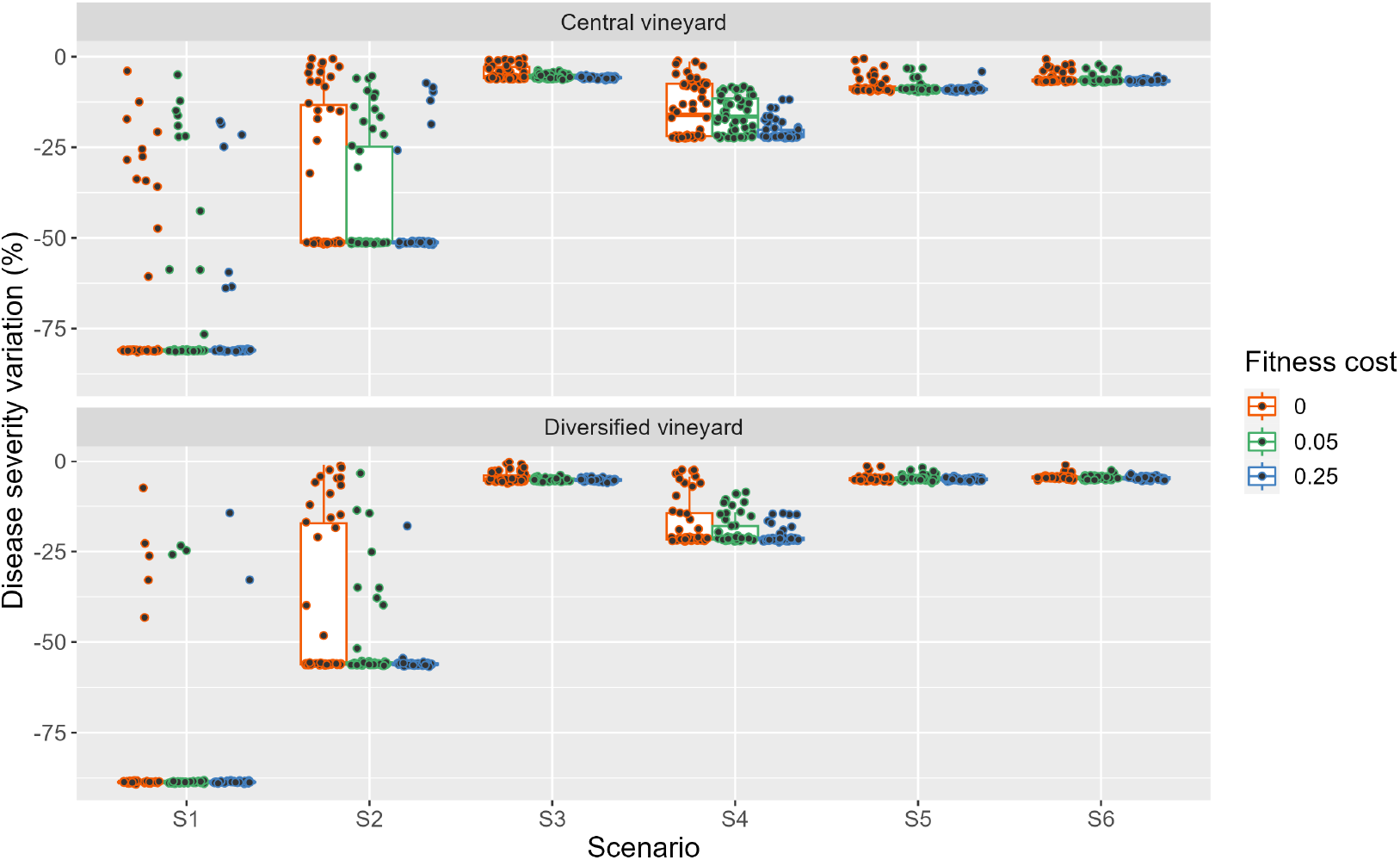
Variability between replicates of epidemiological output across the six scenarios tested. The epidemiological output considered is the percent change in the area under the disease progression curve relative to the baseline scenario S0 (*rAU*). Simulations were performed in the two landscapes of the co-operative territory (“Central” and “Diversified” vineyards) and for three fitness costs. Each boxplot is based on 50 replicates (represented by points) for a given combination of parameters.

**Fig S4.**
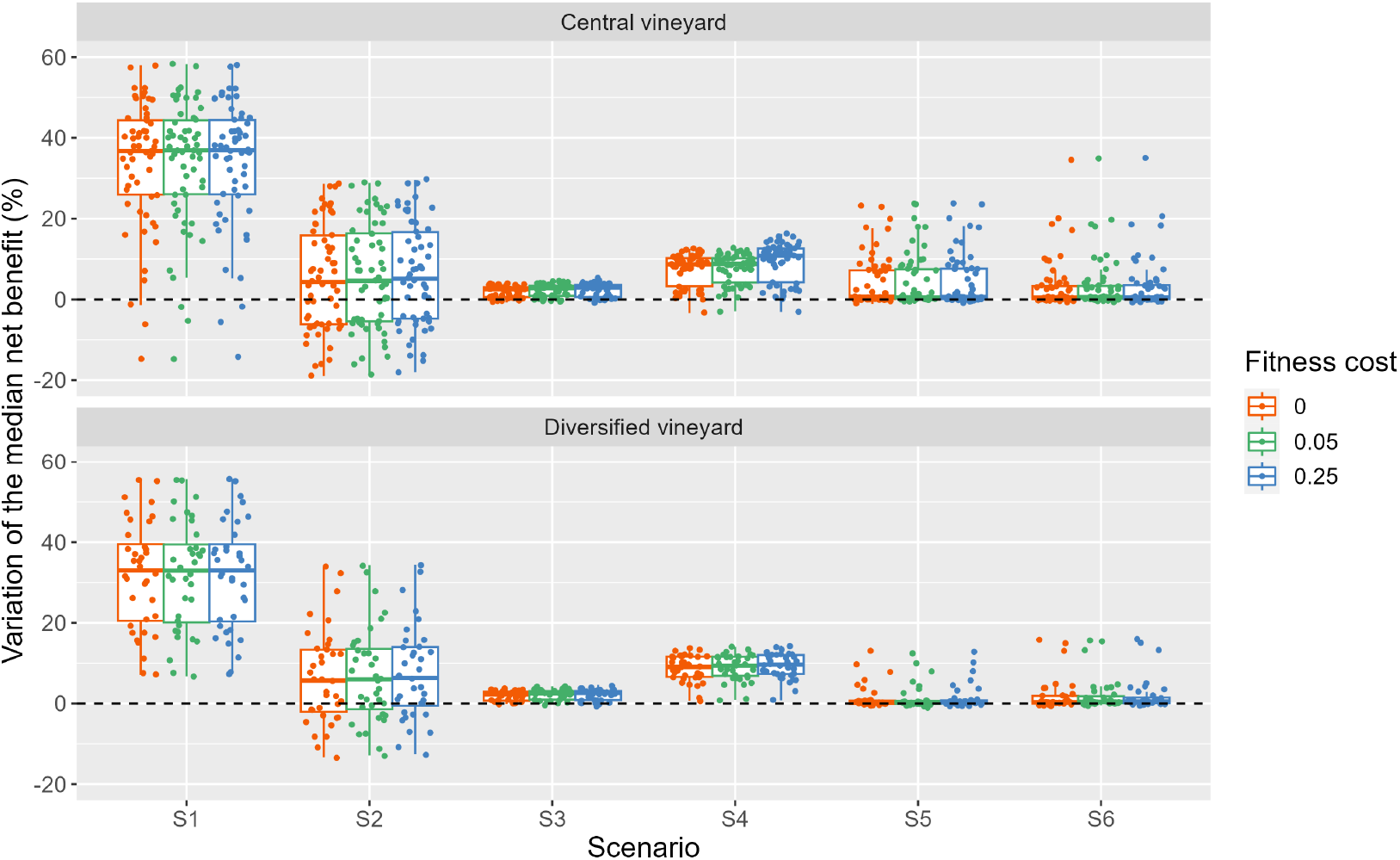
Variability economic output between farmers across the six tested scenarios. For each farmer (57 in the ‘Central’ vineyard and 36 in the ‘Diversified’ vineyard), economic output is expressed as the median (calculated over 50 simulation replicates) percent change in net benefit relative to scenario S0 (*rNB*_*f*_). Results are presented separately for each vineyard (“Central” and “Diversified”) and for three levels of fitness cost. Each boxplot summarizes the distribution of median *rNB*_*f*_ values between individual farmers within each vineyard.

**Fig S5.**
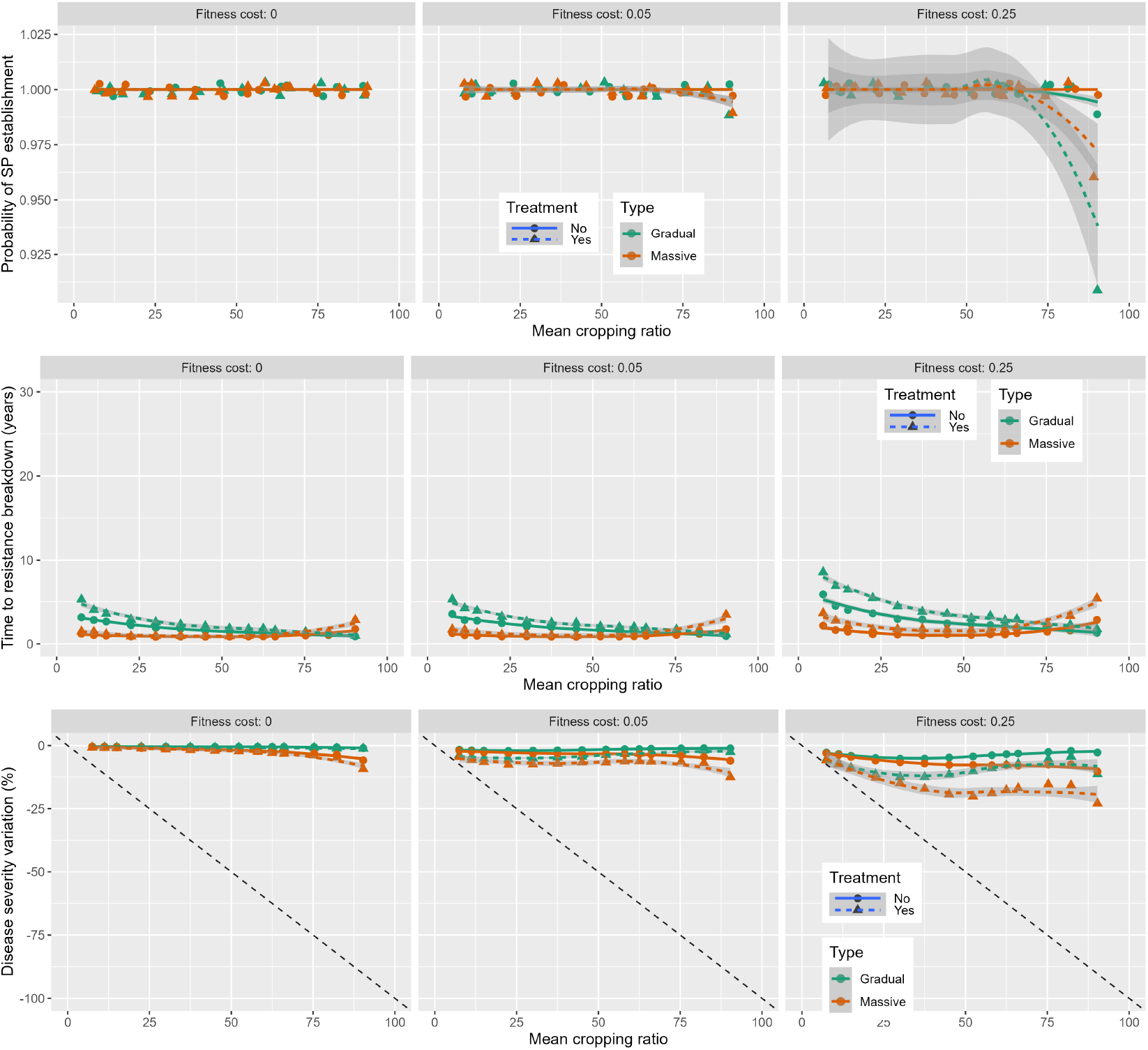
Evolutionary and epidemiological performances of deployment strategies combining massive or progressive RC introduction with the presence or absence of fungicide treatments for a high probability of mutation. Probability of SP establishment (first row), time to establishment given that establishment occurs (second line) and percent change in the area under disease progression curve relative to the baseline strategy S0 (third row) for three fitness costs as a function of the mean percentage of RC cultivated over a period of 30 years. Curves are based on a locally weighted smoothing (loess) method fitted to *p*(*E*_*SP*_), 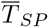 and 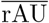 (represented by points). The shaded region represents the 95% confidence interval around the fitted values. The bisector in third row displays the expected variation of disease severity when a RC with perfect resistance impossible to breakdown is deployed.

### Tables

**Table S1.** Probability of pathogen dispersal from fields planted with a susceptible cultivar (SC) to fields planted with a resistant cultivar (RC) in the last year of simulation. This probability is calculated as the probability of a spore propagule produced in an SC field dispersing to a RC field within the landscape, divided by the total probability of the spore dispersing to any field within the landscape. The scenarios (S0 to S6) are defined based on a planting rule, which guides the choice of fields in which resistant cultivars (RC) or susceptible cultivars (SC) are planted. We refer to fields planted more than 30 years before a given year of simulation *y* as “fields ≥ 30 years”. The scenarios concern the “Central” (CV) and “Diversified” (DV) landscapes

| Name Plantation rule | | Probability dispersal SC → RC | |
| --- | --- | --- | --- |
|  |  | CV | DV |
| S0 | SC planted in fields $\geq 30$ years | 0 | 0 |
| S1 | RC planted in fields $\geq 30$ years | 0 | 0 |
| S2 | RC planted in 3.3%/year of the oldest fields | 0.089 | 0.085 |
| S3 | RC planted in fields $\geq 30$ years up to 5% of the area of each farm, otherwise SC planted | 0.016 | 0.014 |
| S4 | RC planted in fields $\geq 30$ years up to 20% of the area of each farm, otherwise SC planted | 0.036 | 0.026 |
| S5 | RC planted in year 1 in fields classified as NTZ Aqua<sup>a</sup>, SC planted in fields $\geq 30$ years | 0.003 | 0.001 |
| S6 | RC planted in year 1 in fields classified as NTZ Riverain<sup>b</sup>, SC planted in fields $\geq 30$ years | 0.004 | 0.002 |

<sup>a</sup> NTZ Aqua: fields in which chemical treatments are restricted due to proximity to water sources;
<sup>b</sup> NTZ Riverain: fields in which chemical treatments are restricted due to proximity to human dwellings (*e.g.* houses, schools).

### S1 Calculating *H*^∗^

*H*^∗^ is the cumulative number of hosts/ha over a cropping season (*T* = 120) in a pure crop and in the absence of disease, calculated analytically with the antiderivative function of Verhust:

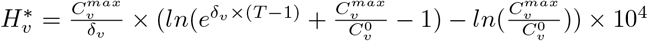

where 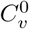 and 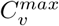 are the plantation and maximal host density of cultivar *v* (in pure crops), respectively, and *δ*_*v*_ is the host growth rate of cultivar *v*. In this work, we assumed that all hosts had the same plantation and maximal densities and growth rates. The *v* index is therefore omitted from the main text.

### S2 Calculating the discount rate applied to the price paid to the grower

The discount rate applied to the price paid to the grower *PriceRed*_*i,y*_ (see eq. 3) was calculated as:

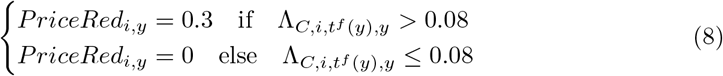

where 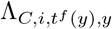 is the severity of infection on a cluster at harvest for field *i* during cropping season *y*. A discount rate of 0.3 is applied to the price paid to the grower if 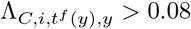 [31, 63]. 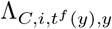 was calculated from the disease severity on leaves, according to a nonlinear model of disease severity progression on clusters described elsewhere [64]:

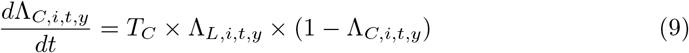

where Λ_*C,i,t,y*_ and Λ_*L,i,t,y*_ represent disease severity on clusters and leaves, respectively, for a given field *i* and a given year *y*. We obtained Λ_*L,i,t,y*_ values by linear interpolation of the terms 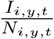 calculated with the model for field *i* and cropping season *y* and at each time-step *t*. The coefficient for transfer from leaves to clusters *T*_*C*_, that is, the increase in disease severity on clusters per unit of disease severity on leaves per day, was set at 0.075 between the “flowering” and “veraison” crop stages (30 ≤ *t* ≤ 107), and was zero otherwise (*t* < 30 or *t* > 107) [64].

